# Subthreshold membrane depolarization powerfully engages intracellular calcium dynamics in the awake mammalian brain

**DOI:** 10.64898/2026.03.05.709685

**Authors:** Yangyang Wang, Hua-an Tseng, Sheng Xiao, Emma Bortz, Yuxin Zhou, Andrew Martin, Heng-Ye Man, Jerome Mertz, Xue Han

## Abstract

Membrane voltage (Vm) regulates spike timing and intracellular signaling. While Vm is extensively modulated by behavior, it is unclear how subthreshold Vm dynamics engage intracellular signaling in the awake mammalian brain. We developed a bicistronic viral vector to express genetically encoded and color compatible voltage and calcium (Ca^2+^) indicators in the same neuron, and simultaneously recorded cellular Vm and Ca^2+^ dynamics in awake mice. We report that prolonged subthreshold Vm depolarization is closely accompanied by prominent large amplitude Ca^2+^ elevation, whereas isolated spikes are coupled with weak Ca^2+^ rise. Additionally, individual spikes differentially engage intracellular Ca^2+^ dynamics depending on post-spiking Vm depolarization, consistent with a prominent role of slow Vm depolarization in regulating cellular signaling. While brief intracranial electrical stimulation consistently leads to Vm depolarization and Ca^2+^ increase, longer stimulation disrupts Vm and Ca^2+^ coupling, highlighting a tightly regulated cellular mechanism that relays slow Vm depolarization to intracellular signaling.

**One-Sentence Summary:** Prolonged membrane depolarization engages cytosolic calcium.

## Introduction

Spiking carries neuronal output and influences intracellular signaling ^1,2^. Spike initiation is tightly regulated by somatic membrane voltage (Vm), which provides a summed measure of a neuron’s intrinsic dynamics and its synaptic inputs ^3^. Spike timing depends not only on the absolute magnitude of Vm depolarization but also its dynamic fluctuations ^3,4^. Typically, spikes are initiated during the rising phase of Vm depolarization and are terminated by rapid Vm repolarization, resulting in a return to the resting condition, a transient hyperpolarization, or a depolarized state known as after-spike-depolarization (ADP) ^3,4^. ADP often triggers subsequent spiking, leading to complex spikes (CSs) characterized by bursts of high frequency spikes superimposed on sustained Vm depolarization ^5^. Beyond transmitting neuronal output, spiking also engages intracellular signaling pathways to modulate synaptic plasticity and neuronal excitability, which influence subsequent spiking responses to synaptic inputs. Among various activity-dependent plasticity mechanisms, cytosolic Ca^2+^ is particularly important. Intracellular Ca^2+^ dynamics are regulated by Vm via voltage-gated Ca^2+^ channels and regulates cellular plasticity through various Ca^2+^-dependent signaling pathways ^1,6^.

Spiking activates high-threshold L- and R-type voltage-gated Ca^2+^ channels that rapidly increase cytosolic Ca^2+^, and thus, transient Ca^2+^ elevation has been anecdotally regarded as a proxy of spiking ^7–11^. With the development of high-performance genetically encoded Ca^2+^ sensors ^7–9^, cytosolic Ca^2+^ dynamics from hundreds or more neurons have been used to characterize neural activities across a wide range of neuroscience studies. In *ex vivo* and *in vitro* neuronal cultures, isolated spikes are generally followed by well-defined Ca^2+^ events characterized by a rapid rising phase that lasts for tens of milliseconds, followed by a slower decay over hundreds of milliseconds or longer ^8,9^. The slow decay phase is influenced by the speed of Ca^2+^ ion dissociation from Ca^2+^ sensors, conformational changes of Ca^2+^ sensors upon Ca^2+^ unbinding, and various cytosolic Ca^2+^ clearance processes. Thus, the rising phase is typically considered to better capture neuronal firing. During bursts of spikes, the cumulative rise in cytosolic Ca^2+^ improves detection sensitivity, leading to the general belief that Ca^2+^ sensors are more sensitive at detecting spike bursts than isolated spikes. Indeed, Ca^2+^ events were found to capture <10% of isolated spikes, but 20-30% of two-spike bursts in the brain ^12^.

Spiking-mediated Ca^2+^ influx through the plasma membrane influences Vm and membrane excitability ^13–15^. For example, patch clamp studies in brain slices have demonstrated that increases in cytosolic Ca^2+^ are critical for prolonged somatic Vm depolarization during CSs^16–18^. Blocking L-type Ca^2+^ channels in behaving mice reduced such depolarizations^19,20^. The prominent and long-lasting somatic Vm depolarization following spiking can recruit diverse voltage-gated Ca^2+^, Na^+^ and K^+^ channels, as well as Ca^2+^-conducting NMDA receptors ^21–26^, further shaping Vm and Ca^2+^ dynamics. In addition to spiking-related Ca^2+^ entry from the plasma membrane, cytosolic Ca^2+^ is also regulated by low-threshold voltage-gated T-type Ca^2+^ channels, Ca^2+^ release from intracellular stores, controlled Ca^2+^ clearance, and other cellular processes ^27,28^.

Over the years, *in vitro* studies in brain slices and cultured neurons have demonstrated a close relationship between Vm and cytosolic Ca^2+ 29,30^, which depend on voltage-gated Ca^2+^ channels along with many other ion channels, including persistent Na^+ 31,32^, K^+ 5,16,26^, and HCN channels ^23^. However, in behaving mammals, neurons receive extensive and highly dynamic synaptic inputs, resulting in more substantial subthreshold Vm fluctuations and higher conductance than neurons *in vitro*, where more inputs are severed or inactive ^33^. Furthermore, depending on the intrinsic biophysical properties, Vm in some neurons could be preferentially engaged by inputs at certain frequencies, which may differentially influence cytosolic Ca^2+ 22,28,34^. Thus, in the brains of awake mammals, the relationship between Ca^2+^ and Vm is likely much more complex ^10,12^.

To determine how a neuron’s subthreshold Vm and spiking influence cytosolic Ca^2+^ in the awake mammalian brain, ideally, one would record both simultaneously in the same cell. However, it is difficult to obtain sufficient expression of multiple activity sensors ^35^. Accordingly, we generated bicistronic AAV viral vectors containing the high performance voltage sensor SomArchon and the color-compatible Ca^2+^ sensor GCaMP7f or GCaMP8m ^8,9,36,37^. Both GCaMP7f and 8m exhibit excellent sensitivity to increases in cytosolic Ca^2+^, but with different Ca^2+^ binding affinities (Kd of 150 nM for GCaMP7f and 108 nM for GCaMP8m). Additionally, GCaMP8m has a faster onset time of 7.4 ms compared to 26.8 ms for GCaMP7f, whereas GCaMP7f has a brighter basal fluorescence that may be advantageous for identifying transduced neurons ^8,9^. Using these AAV vectors, we simultaneously measured somatic Vm and Ca^2+^ in cultured rat neurons, human iPSC-derived neurons, and in neurons across the striatum, CA1 and visual cortex in awake mice, and accessed the effects of intracranial stimulation.

### Color compatible SomArchon and GCaMP7f enable simultaneous cellular voltage (Vm) and Ca^2+^ imaging in rat primary neuron cultures and human iPSC-derived neuron cultures

To examine the relationship between somatic Vm and cytosolic Ca^2+^ within the same neuron, we first designed a bicistronic AAV vector linking the near-infrared, soma-targeted SomArchon voltage sensor and the color-compatible green GCaMP7f Ca^2+^ sensor with the self-cleaving P2A peptide (AAV9-Syn-SomArchon-P2A-GCaMP7f, abbreviated as SomArchon-GCaMP7, Fig. 1&2 and Fig. S1). Transducing rat hippocampal neuron cultures (Fig. 1B-E) and human iPSC-derived neuron cultures (Fig. 1F-I) with SomArchon-GCaMP7f revealed expected membrane-targeted SomArchon expression and cytosolic GCaMP7f expression (Fig. 1B&F).

**Figure 1.**
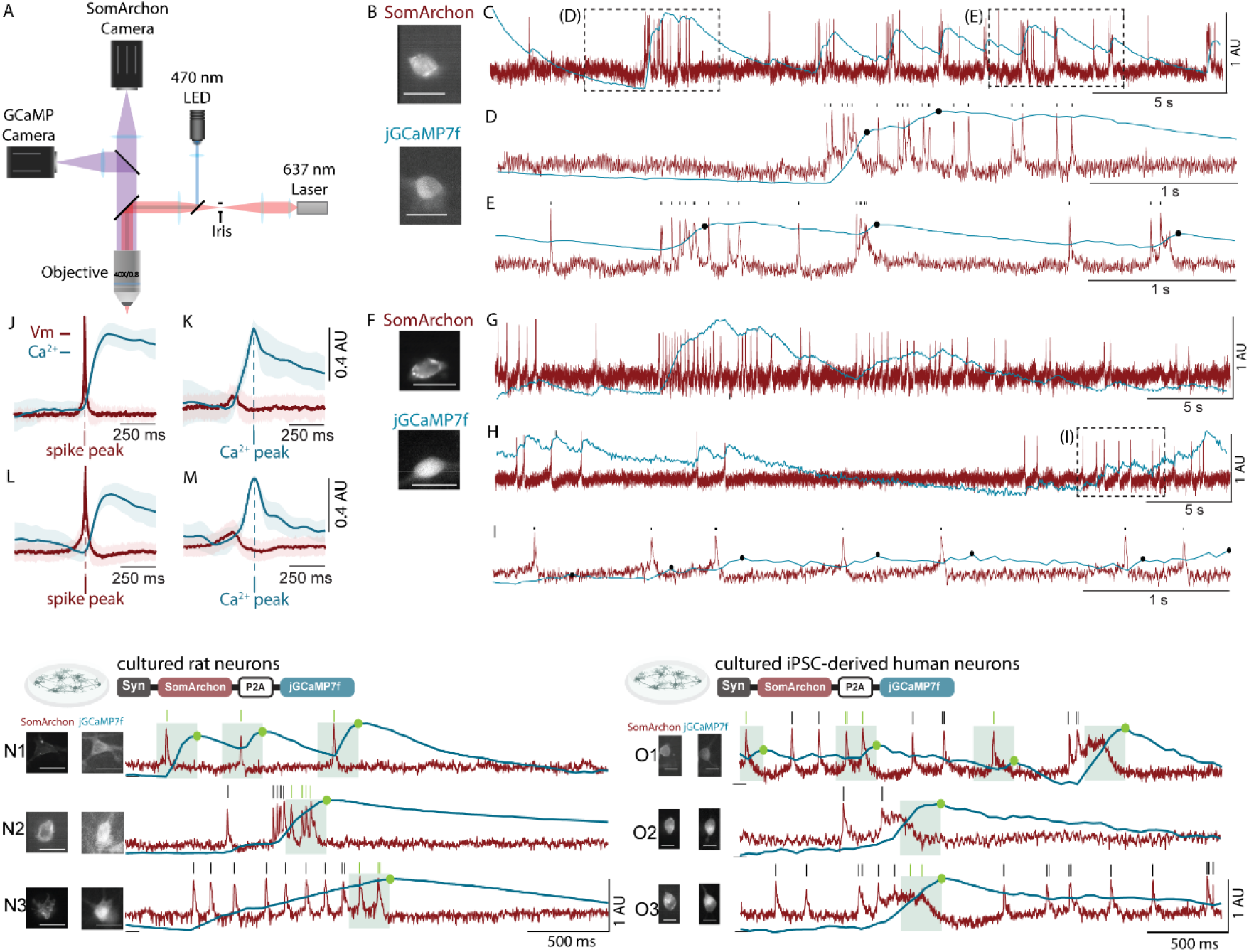
Simultaneous recording of Vm and Ca^2+^ in cultured rat neurons and human iPSC-derived neurons. (**A**) Schematic of the custom microscope for simultaneous recording of SomArchon and GCaMP fluorescence (details in Fig. S1). (**B**) An example cultured rat neuron transduced by AAV9-Syn-SomArchon-P2A-GCaMP7f (SomArchon-GCaMP7f). Top, the maximum-minus-minimum projection image of SomArchon fluorescence; bottom, corresponding GCaMP7f fluorescence. Scale bars: 30 µm. (**C**) Vm (red) and Ca^2+^ (blue) traces simultaneously recorded from the neuron shown in (B). (**D, E**) Zoomed in of the trace shown in (C). Black ticks: spikes, black dots: Ca^2+^ event peaks. (**F-I**) Similar to (B-E), but for an example cultured human iPSC-derived neuron. (**J**) Population Vm (red) and Ca^2+^ (blue) dynamics aligned to spike peaks across rat neurons (n=11). Shade area: standard deviation. (**K**) Population Vm (red) and Ca^2+^ (blue) dynamics aligned to Ca^2+^ event peaks across rat neurons (n=11). Shade area: standard deviation. (**L, M**) Similar to (**J, K**), but for cultured human neurons (n=16). (**N1-N3**) Additional example recordings of Vm (red line) and Ca^2+^ (blue line) from three rat cultured neurons expressing SomArchon-GCaMP7f, showing the spikes within (green tick) versus outside (black tick) of the 250 ms window prior to Ca^2+^ event peaks (green shading). Green dots: Ca^2+^ event peaks. Scale bars: 30 µm. (**O1-O3**) Similar to (N1-N3), but for three human iPSC-derived neurons. Scale bars: 15 µm. 1 AU=100% dynamic range.

After confirming robust expression of SomArchon and GCaMP7f in the same cell, we simultaneously recorded their fluorescence using a custom wide-field microscope containing a SomArchon light path and a GCaMP light path (Fig. 1A, Fig. S1, and Methods). SomArchon fluorescence was collected at 500 Hz for rat culture and 667 Hz for human culture, and GCaMP7f fluorescence at 20 Hz. Each field of view was imaged for 30 or 45 seconds per trial, for up to 5 trials, with an intertrial interval of ∼30 seconds. The corresponding image frames were motion corrected, and individual neuron soma manually segmented. SomArchon and GCaMP7f fluorescence across all pixels within a neuron was then averaged, background subtracted, detrended and normalized to obtain Vm and Ca^2+^ traces respectively (see Methods). Due to the slower sampling rate of GCaMP7f, Ca^2+^ traces were upsampled to match the corresponding Vm traces. Ca^2+^ and Vm traces were then scaled between 0 and 1 (from minimum to maximum fluorescence, 1 AU = 100%) for all subsequent analysis.

We first quantified the relationship between spiking and Ca^2+^ elevation. Individual spikes were identified as rapid, large amplitude Vm depolarizations, and Ca^2+^ events were identified as significant increases in Ca^2+^ traces (see Methods, and Fig. 1D, E&I). Consistent with previous *in vitro* observations ^29,38^, we found that in both rat and human neuron cultures, individual spikes were accompanied by consistent, but small Ca^2+^ rises followed by a slow decay (Fig. 1C, G&H). Clusters of spikes were generally accompanied by larger Ca^2+^ rises, and often with a visibly graded increase accompanying each spike within a cluster (Fig. 1C-I and N-O). Aligning Ca^2+^ traces to individual spikes revealed that Ca^2+^ peaked at 264±135 ms and 186±45 ms after spikes in rat and human neurons respectively (Fig. 1J&L). Similarly, aligning Vm to Ca^2+^ event peaks showed that Vm depolarization proceeded Ca^2+^ peaks by 152±36 ms and 145±93 ms in rat and human cultures respectively (Fig. 1K&M).

As Ca^2+^ events have often been considered a surrogate of neuronal firing ^11,39,40^, we estimated the fraction of spikes captured by the rising phase of Ca^2+^ events, and found that ∼70% of spikes occurred within 250 ms (green shadings in Fig. 1N&O) prior to Ca^2+^ event peaks (rat neurons: 72±19%, n=11; human neurons: 67±17%, n=16). Thus, neuronal firing and cytosolic Ca^2+^ events are temporally associated *in vitro*, and SomArchon and GCaMP7f allow for simultaneous recording of Vm and Ca^2+^ in cultured rat neurons and human iPSC-derived neurons, providing an easily scalable tool for large scale simultaneous Ca^2+^ and Vm analysis.

### Simultaneous imaging of Vm and Ca^2+^ in the same neurons in awake mice reveals prominent Ca^2+^ rise during prolonged Vm depolarization in CA1, dorsal striatum and visual cortex

After confirming the ability to simultaneously record Vm and Ca^2+^ in cultured neurons, we evaluated SomArchon-GCaMP7f performance in the hippocampal CA1 of awake mice. Briefly, we surgically implanted an optical window coupled with an infusion cannula over the CA1, along with a headplate for awake head-fixed imaging. Approximately one-week post-surgery, AAV9-Syn-SomArchon-P2A-GCaMP7f was infused into the CA1 through the cannula, which mediated robust expression of SomArchon and GCaMP about two weeks after viral infusion (Fig. 2B, H, L&P).

**Figure 2.**
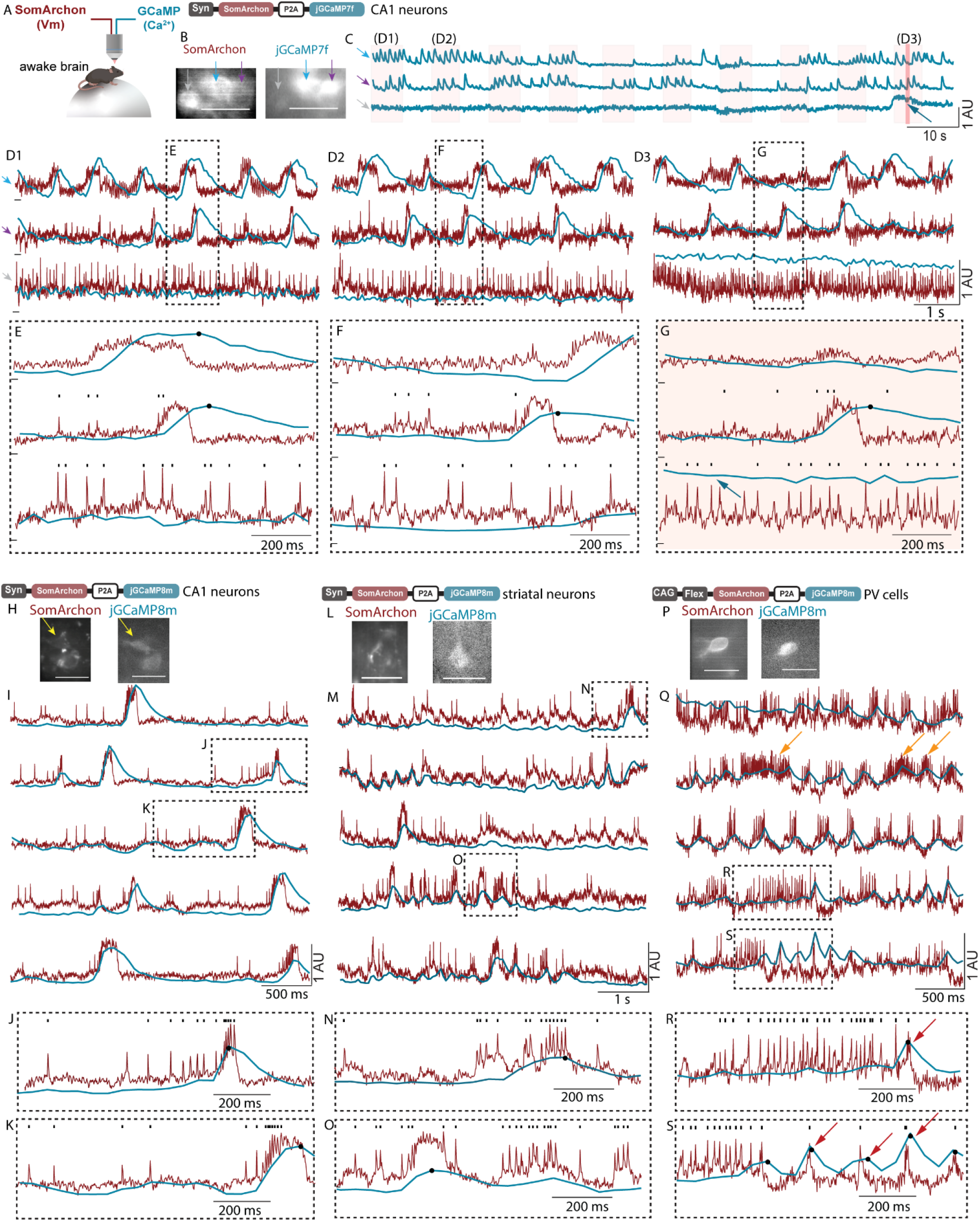
Simultaneous Vm and Ca^2+^ recordings from the same neurons in CA1, striatum, and visual cortex in awake mice. (**A**) Schematics of simultaneous Vm and Ca^2+^ imaging in awake mice. (**B**) The maximum-minus-minimum projection image of SomArchon fluorescence and corresponding GCaMP7f fluorescence in an example CA1 recording containing three neurons. Scale bars: 30 µm. (**C**) Simultaneously recorded Ca^2+^ traces from the three neurons shown in (B), with the pink shadings indicating the periods when SomArchon fluorescence was recorded. (**D1-D3**) Example Vm (red) and Ca^2+^ (blue) traces during the periods highlighted in (C). Colored arrows on the left indicate the corresponding neurons marked in (B). (**E-G**) Zoom-in view of the periods highlighted by dotted boxes in (D). Black ticks: spikes; Black dots: Ca^2+^ event peaks; blue arrows in C and G highlight a slow but prominent Ca^2+^ elevation observed in the gray neuron. (**H-K**) Similar to (B-G), but for an example CA1 neuron expressing SomArchon-GCaMP8m. Shown are five periods of simultaneous Vm and Ca^2+^ recordings from the neuron shown in (H). (**J, K**) Zoom-in view of the periods highlighted by dotted boxes in (I). Black ticks: spikes; Black dots: Ca^2+^ event peaks. (**L-O**) Similar to (H-K), but for an example dorsal striatal neuron expressing SomArchon-GCaMP8m. (**P-S**) Similar to (H-K), but for a visual cortex PV cell expressing SomArchon-GCaMP8m. Orange arrows in (Q) exemplify tonic firing periods, and red arrows in (R-S) exemplify Ca^2+^ events detected during burst firing periods. 1 AU=100% dynamic range.

We then imaged GCaMP at 20 Hz continuously for 1-2 minutes, and SomArchon at 500 Hz intermittently for 3-6 seconds every 6-12 seconds (Fig. 2C), while mice were head-fixed and voluntarily locomoting (Fig. 2A and Fig. S1). To avoid signal crosstalk, we recorded from well isolated individual neurons (Fig. S2). The recorded SomArchon and GCaMP fluorescence traces were extracted for each neuron, visually inspected to ensure the signal quality, background subtracted, detrended and normalized to maximum fluorescence intensity (see Methods). Spikes and Ca^2+^ events were subsequently identified in the corresponding Vm and Ca^2+^ traces. The spike rate and Ca^2+^ event rate measured with these sensors are in general agreement with previous studies using SomArchon and GCaMP7f alone^3,8,41^ (Table S1).

Different from the rather stable subthreshold Vm *in vitro*, we observed frequent and prominent slow Vm depolarization in awake mice (Fig. 2D). Strikingly, prominent slow Vm depolarization was consistently accompanied by large-amplitude Ca^2+^ rises (Fig. 2B-G). In contrast, individual spikes not accompanied with slow Vm depolarization, particularly during rapid firing, were generally associated with small amplitude Ca^2+^ fluctuations, suggesting that subthreshold slow Vm dynamics play an important role in regulating cytosolic Ca^2+^.

As GCaMP7f and 8m exhibit distinct Ca^2+^ binding affinities, kinetics, and brightness ^8,9^, we additionally generated the bicistronic AAV vector AAV9-Syn-SomArchon-P2A-GCaMP8m (SomArchon-GCaMP8m) to express SomArchon and GCaMP8m nonspecifically in neurons, and AAV9-CAG-Flex-SomArchon-P2A-GCaMP8m to express them in Cre-positive cells. Further testing of SomArchon-GCaMP8m in CA1 (Fig. 2H-K and Fig. S3A) and dorsal striatum neurons (Fig. 2L-O and Fig. S3B) and visual cortex parvalbumin-positive interneurons (PV cells) in PV-Cre mice (Fig. 2P-S) revealed similar prolonged Vm depolarization that was accompanied by large amplitude Ca^2+^ events. In particular, GCaMP signals exhibited little fluctuations following individual spikes in tonically fast firing PV cortical neurons (Fig. 2R-S). However, large GCaMP events emerged when PV cells transitioned to burst firing (Fig. 2R-S), consistent with the generally low event rate measured with Ca^2+^ indicators in PV cells ^42,43^. Similarly, fast spiking neurons in the striatum and CA1 also showed slow Ca^2+^ changes (Fig. 2B-G, and Fig. S3A, S3B).

### Ca^2+^ event rising phase captures transient increase in spike rate, even though it is not reliable with detecting individual spikes

As most Ca^2+^ events *in vivo* exhibit the characteristic fast rising and slow decay, we identified the rising phase of each Ca^2+^ event (Fig. 3A, C, E). In CA1, 26±12% (n=31 neurons) of spikes occurred on the rising phase of GCaMP7f (Fig. 3A), and the spike rate during the rising phase was significantly higher than during the non-rising phase (Fig. 3B). Like GCaMP7f, 30±16% (n=36 neurons) of spikes occurred on the GCaMP8m rising phase (Fig. 3C), leading to a significantly higher firing rate during the rising phase (Fig. 3D). Similarly, in the striatum, 27±11% (n=21 neurons) of spikes occurred on the GCaMP8m rising phase, and the firing rate is higher during rising phase (Fig. 3E, F). Thus, Ca^2+^ event rising phase captures transient spike rate increase in awake mice, even though it is not reliable with detecting individual spikes.

**Figure 3.**
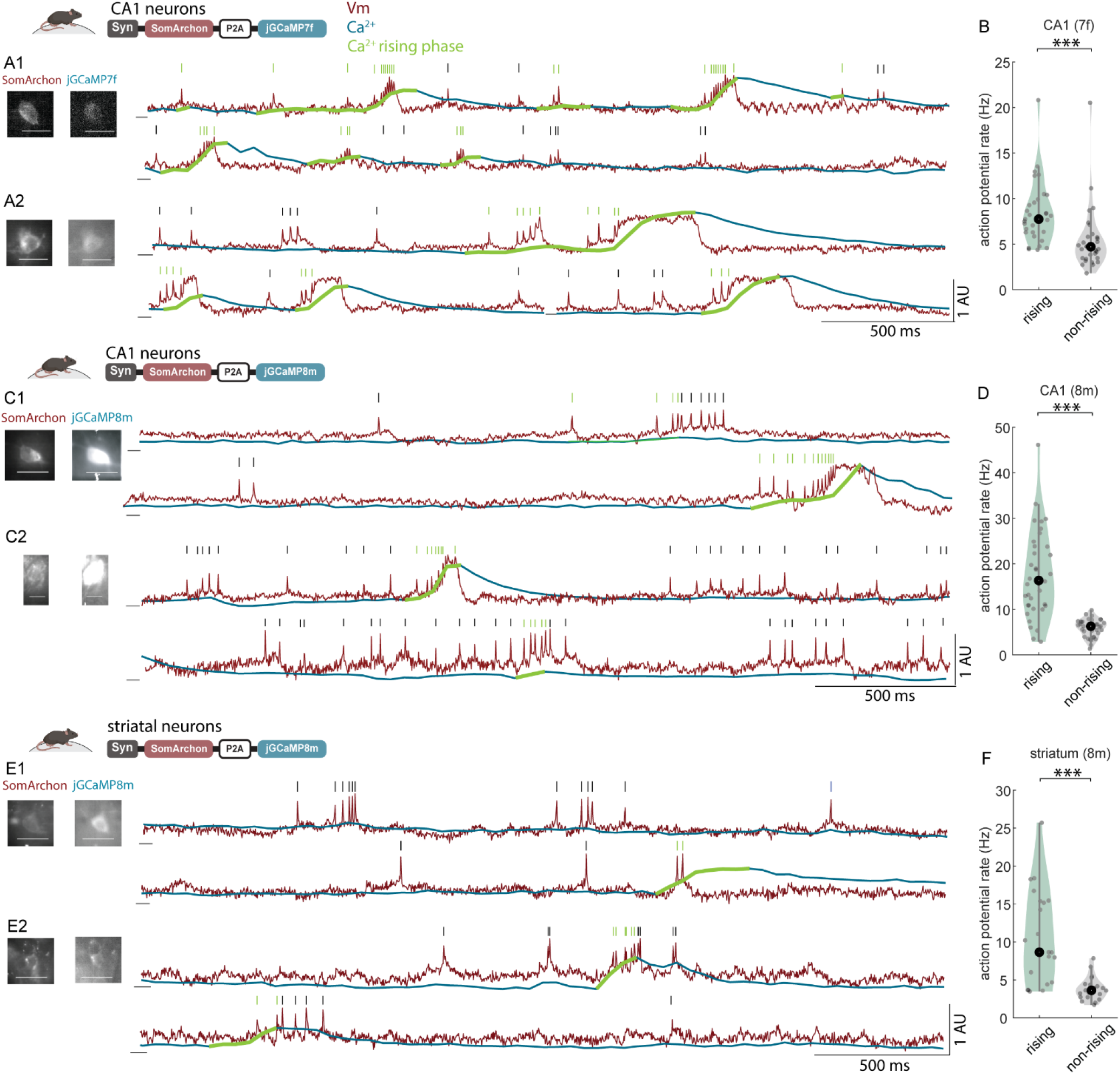
Ca^2+^ event rising phase was accompanied by elevated firing rate, though failed to capture many spikes. (**A1-A2**) Example recordings of Vm (red line) and Ca^2+^ (blue line) from two CA1 neurons in awake mice expressing SomArchon-GCaMP7, showing spikes within (green tick) versus outside (black tick) the Ca^2+^ rising phase (green line overlaid on the blue trace). Scale bar: 30 µm. (**B**) Firing rate during rising and non-rising phase of Ca^2+^ events in CA1 neurons expressing SomArchon-GCaMP7. Wilcoxon rank sum test, p=2.4e-5. (**C-D**) Similar to (A-B), but for CA1 neurons expressing SomArchon-GCaMP8. p=2.3e-9. Scale bar: 30 µm in (C1) and 15 µm in (C2). (**E-F**) Similar to (A-B), but for striatal neurons labelled with SomArchon-GCaMP8. p=3e-5. Scale bar: 30 µm. 1 AU=100% dynamic range.

### Spikes accompanied by Vm depolarization, particularly during complex spikes (CS), are better coupled to cytosolic Ca^2+^ than those without Vm depolarization

As GCaMP8m is slightly more sensitive with detecting spikes, though not significantly different from GCaMP7f, subsequent analysis was focused on SomArchon-GCaMP8m. Spikes are sometimes accompanied by after spike depolarization (ADP ^44^), which creates a period of heighted excitability and permits subsequent high frequency firing. The co-occurrence of ADP and fast spiking are traditionally referred to as complex spikes (CSs), often captured as bursts of spikes with short inter-spike-intervals (ISIs) in extracellular recordings that cannot detect Vm. Being able to detect both spiking and ADPs with SomArchon, we classified spikes based on ISIs and ADPs. Specifically, single spikes (SSs) were defined as well isolated spikes with ISIs greater than 30 ms. SSs were further divided into SSs with ADP (SS-w-ADP) if the mean Vm amplitude during 5-30 ms post spiking was higher than pre-spiking, and SSs without ADP (SS-wo-ADP) if otherwise (Fig. 4A-C black and gray ticks, see Methods). CSs were defined as a continuous sequence of spikes with ISIs shorter than 30 ms and with ADPs, and the onset of CS was defined as the first spike within the sequence (Fig. 4A-C green and orange ticks, see Methods).

**Figure 4.**
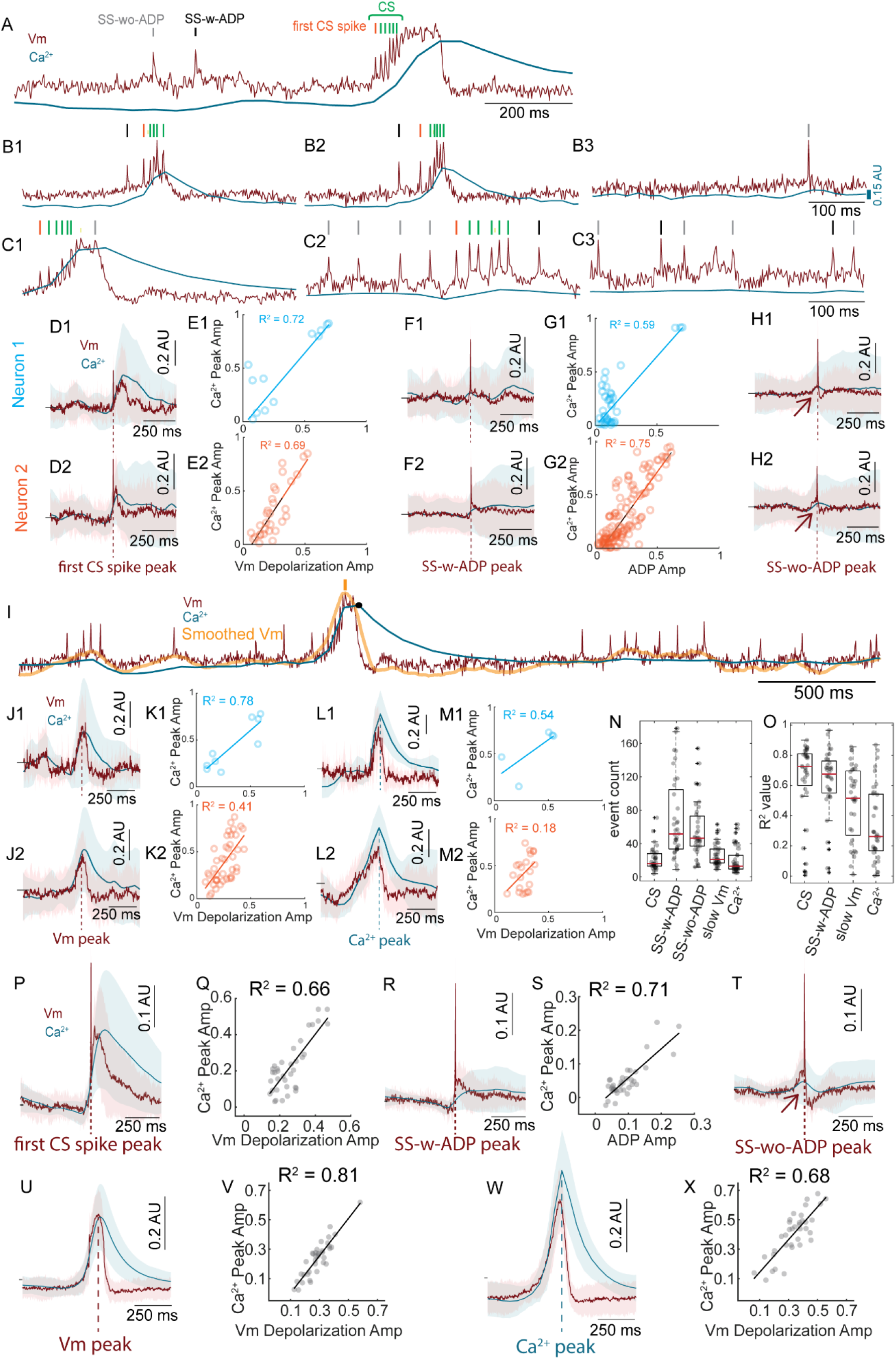
Slow subthreshold Vm depolarization is tightly correlated with cytosolic Ca^2+^ elevation in CA1 neurons. (**A**) An example recording from a CA1 neuron expressing SomArchon-GCaMP8. Vm trace (red) was overlaid with Ca^2+^ trace (blue). The colored ticks indicate the identified single spikes without ADP (SS-wo-ADP, gray), single spikes with ADP (SS-w-ADP, black), and complex spike (CS, first spike in orange and subsequent spikes in green). (**B1-B3**) Additional example recordings from another CA1 neuron. (**C1-C3**) Similar to (B1-B3), but from another CA1 neuron. (**D1**) Vm (red) and Ca^2+^ (blue) dynamics aligned to CS onset from this neuron. Solid lines: mean. Shade area: standard deviation (n=16 events). (**D2**) Similar to (D1), but for another example neuron, (n=33 CS events). (**E**) Vm depolarization peak amplitude versus Ca^2+^ peak amplitude across all CS events for the two neurons, respectively. Each color corresponds to each neuron. Each circle represents each CS event. Solid lines indicate linear regression, with the R^2^ values indicated. (**F-G**) Similar to (D-E), but for SS-w-ADP (F1: n=45 events; F2: n=152 events). (**H**) Similar to (F), but for SS-wo-ADP (H1: n=84 events; H2: n=90 events). (**I**) Example Vm traces (red), overlaid with smoothed subthreshold Vm (orange) and Ca^2+^ traces (blue), in one example neuron expressing SomArchon-GCaMP8. The orange ticks indicate slow Vm depolarization peaks, and the black dots indicate Ca^2+^ peaks. (**J1-J2**) Similar to (D1-D2), but for Vm and Ca^2+^ dynamics aligned to slow Vm depolarization peaks. (J1: n=9 slow Vm events; J2: n=41 slow Vm events). (**K**) Vm depolarization peak amplitude versus Ca^2+^ peak amplitude across all slow Vm events for the two neurons shown in (D1, D2), respectively. Each color corresponds to each neuron. Each circle represents each CS event. Solid lines indicate linear regression, with the R^2^ values indicated. (**L-M**), Similar to (J-K), but for Ca^2+^ peaks. (L1: n=5 Ca^2+^ events; L2: n=21 Ca^2+^ events). (**N**) Boxplots of the event counts for each neuronal activity category (CS, SS-w-ADP, SS-wo-ADP, slow Vm, and Ca^2+^ peaks) (n=36 neurons). Each dot corresponds to a neuron and + marks outliers. The red line is the median, and whiskers indicate the minimum and maximum. (**O**) Similar to (N), but for the R^2^ values of the linear regression between Vm and Ca^2+^ amplitudes for four neuronal activity categories (CS, SS-w-ADP, slow Vm, and Ca^2+^ peaks). (**P**) Population Vm (red) and Ca^2+^ (blue) dynamics aligned to CS onset from SomArchon-GCaMP8 expressing CA1 neurons. Solid lines: mean. Shade area: standard deviation (n=36 neurons). (**Q**) Vm depolarization peak amplitude versus Ca^2+^ peak amplitude across neurons shown in (P). Each dot corresponds to one neuron. Black line indicates linear regression, with the R^2^ values indicated. (**R, S**) Similar to (P, Q), but for SS-w-ADP. (**T**) Similar to (R), but for SS-wo-ADP. (**U**) Population Vm and Ca^2+^ dynamics aligned to slow Vm depolarization peaks across neurons expressing SomArchon-GCaMP8 (n=36 neurons). Shade area: standard deviation. (**V**) Vm depolarization peak amplitude versus Ca^2+^ peak amplitude across neurons. (**W, X**) Similar to (U, V), but for Vm and Ca^2+^ dynamics aligned to Ca^2+^ event peaks. 1 AU=100% dynamic range.

Across the recorded CA1 neurons, 28±16% of spikes were part of a CS, 33±12% were SS-w-ADP, and 28±9% were SS-wo-ADP. The observed CS occurrence is in general agreement with previous studies using patch clamp ^45^ and voltage imaging ^3^, and extracellular recordings that approximated CSs as spike bursts due to the inability to record Vm ^3,21,46–48^. Additionally, we observed that many neurons switched between SSs and CSs (Fig. S3C), consistent with our previous study using SomArchon alone ^3^.

For each neuron, we aligned both Vm and Ca^2+^ traces to each CS onset, the peak of the first spike within a CS and characterized the relationship of their peak amplitude (Fig. 4D). Specifically, we first smoothed the Vm trace by a 50-ms time window, to match the sampling rate of the GCaMP recordings, and then identified the peaks of Vm and Ca^2+^ traces within 5-500 ms post CS. Interesting, we noted a strong linear correlation between these two peaks, suggesting a close coupling of CS and intracellular Ca^2+^ rise at the individual neuron level (Fig. 4E, N&O). Across neurons, similar analysis revealed concomitant large amplitude Vm depolarization (25.9±9.2% of the maximum-minimum Vm dynamic range, mean±std, n=36) and Ca^2+^ rise (22.2±14.5% of the maximum-minimum Ca^2+^ dynamic range, mean±std, n=36, Fig. 4P). Vm depolarization peaked at ∼57 ms after CS onset, consistent with the gradual ramping of depolarization during CS (Fig. 4P). The peak amplitude of Vm depolarization versus Ca^2+^ elevation showed a strong linear relationship across neurons (Fig. 4Q), consistent with the single neuron analysis (Fig. 4O). To rule out potential contribution of fluorescence signals from adjacent neurons within the same field of view, we aligned Vm and Ca2+ traces of one neuron to CS onsets of another simultaneously recorded neighboring neuron and detected no obvious Vm or Ca^2+^ change, confirming a lack of signal cross contamination (Fig. S7). Thus, CS robustly engages intracellular Ca^2+^ in awake mice, consistent with previous observation using simultaneous intracellular recordings and Ca^2+^ imaging in anesthetized animals.

While less prominent than CS, we also observed a small but significant Ca^2+^ peak following SS-w-ADP at both single neuron and population levels (Fig. 4F, R, 5.14±5.38% of the maximum-minimum Ca^2+^ dynamic range, mean±std, n=36), and ADP and Ca^2+^ peak amplitude also exhibited a positive correlation (Fig. 4G, O&S). In contrast, following SS-wo-ADP, Ca^2+^ increase was negligible (Fig. 4H&T).

**Figure 5.**
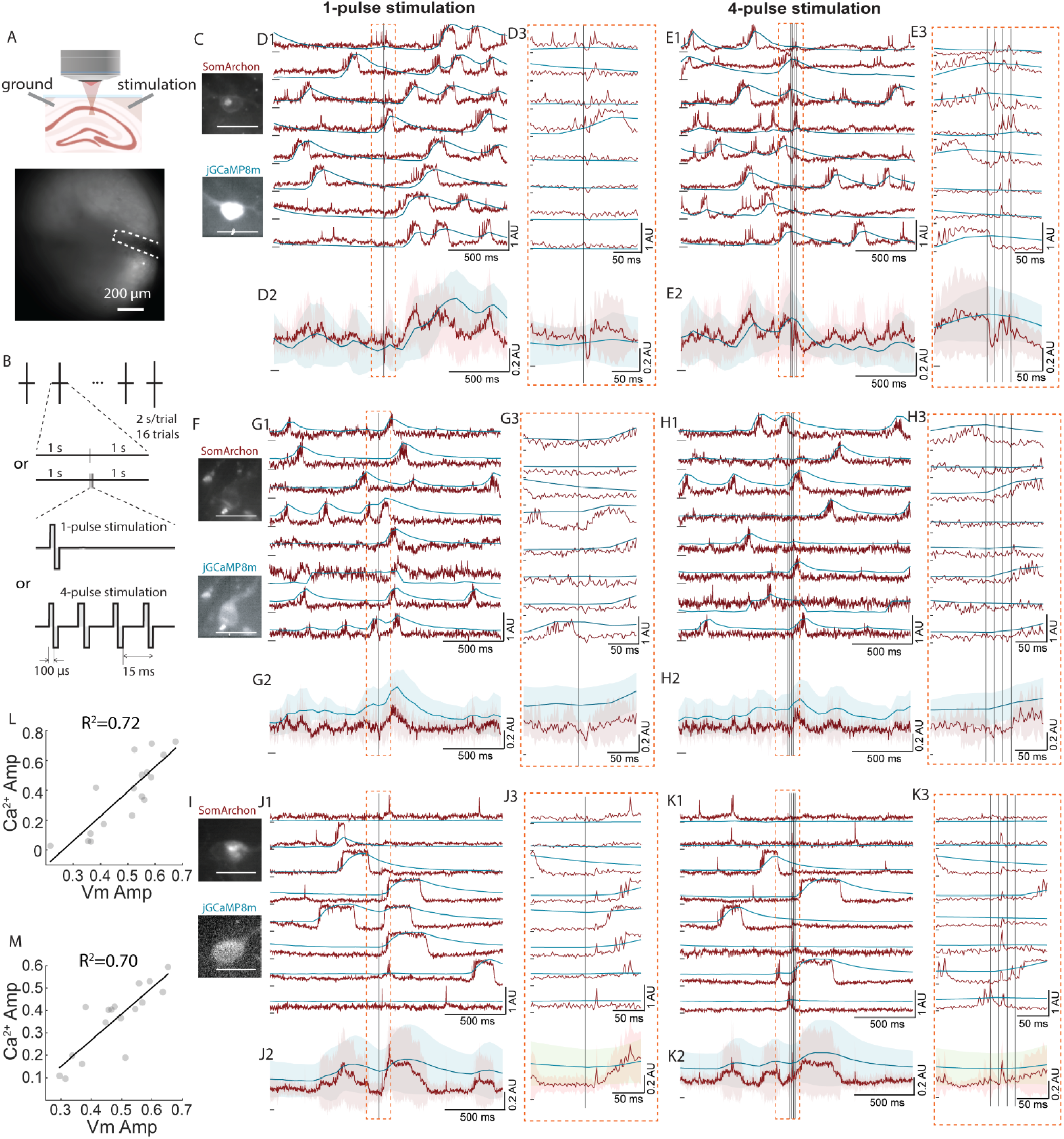
Brief intracranial electrical stimulation evokes consistent Vm depolarization and cytosolic Ca^2+^ increase in CA1 neurons in awake mice. (**A**) Top: schematic of the experimental setup, illustrating the relative position of the imaging field of view (FOV) to the electrode pair. Bottom: an example imaging FOV with a nearby electrode highlighted. (**B**) One or four pulse electrical stimulation protocols. Each trial was 2 s, with stimulation onset at 1 s. Each pulse was 100 µs/phase, and the inter-pulse interval was 15 ms for four pulse trials. (**C**) The maximum-minus-minimum projection image of SomArchon fluorescence and corresponding GCaMP8m fluorescence in an example CA1 neuron. Scale bars: 30 µm. (**D1**) Vm (red) and Ca^2+^ (blue) traces across individual 1-pulse trials for the neuron shown in (C). Black lines: onset of each electrical pulse. (**D2**) Mean evoked response across trials shown in (D1). Shaded areas are standard deviations. (**D3**) Zoomed in view of the 200 ms time window around the pulse onset, highlighted by the orange box in (D1-D2). (**E1-E3**) Similar to (D1-D3), but for 4-pulse stimulation trials from the same neuron in (C). (**F-H**) and (**I-K**) Similar to (C-E), but for two other example CA1 neurons. (**L**) Scatter plots of the peak amplitudes of evoked Vm versus Ca^2+^ within 250 ms after the onset of 1-pulse stimulation (n=17 neurons). Black line indicates linear regression, with the R^2^ value indicated. (**M**) Similar to (L), but for the same 17 neurons during 4-pulse stimulation. 1 AU=100% dynamic range.

Similar analysis in dorsal striatal neurons with SomArchon-GCaMP8 revealed identical results as in CA1, even though the amplitudes of Vm depolarization and the corresponding Ca^2+^ event were generally less correlated than CA1 (Fig. S6A-E). Finally, recordings made with SomArchon-GCaMP7 further corroborated the observed Vm and Ca^2+^ relationship (Fig. S5A-E). Thus, spikes followed by ADPs robustly elevate cytosolic Ca^2+^, particularly during CS, whereas spikes without ADP exert much weaker influence on intracellular Ca^2+^ dynamics. These results highlight a prominent role of ADP on engaging cytosolic Ca^2+^, suggesting that different spiking modes engage distinct cellular Ca^2+^ dynamics.

### Slow subthreshold Vm depolarization is tightly correlated with cytosolic Ca^2+^ elevation in both the CA1 and striatum

To further investigate the relationship between slow subthreshold Vm depolarization and Ca^2+^ elevation, we removed the impact of spiking by smoothing Vm traces using a 100.6 ms moving window, essentially computing subthreshold slow dynamics below 10 Hz. We then identified the peaks of slow Vm depolarization (Fig. 4I), and aligned Ca^2+^ traces to these peaks in each neuron (Fig. 4J, K&O) across all neurons. In CA1 neurons, slow Vm depolarization is consistently accompanied by Ca^2+^ increase (Fig. 4U), with a lag of 38±66 ms (n=36). Similarly, aligning slow Vm traces to Ca^2+^ event peaks revealed similar results across Ca^2+^ events in each neuron (Fig. 4L, M&O). Across all neurons Vm depolarization preceded Ca^2+^ event peaks by 19±18 ms (n=36, Fig. 4W). Again, the peak amplitude of Ca^2+^ was linearly correlated with that of Vm depolarization (Fig. 4V&X), in line with the observations from CS and SS-w-ADP. The same analysis in striatal neurons revealed consistent results (Fig. S6F-I).

To examine the potential contribution of SomArchon and GCaMP basal protein expression levels in the observed correlation between their peak amplitudes, we compared both indicators’ raw fluorescence and peak amplitudes in CA1 (Fig. S8A-F) and striatum (Fig. S8G-I), and detected little correlations in all comparisons. Similar findings were also obtained with SomArchon-GCaMP7f (Fig. S5F-I). Thus, slow Vm depolarization is tightly coupled to cytosolic Ca2+ elevation, further supporting a role of subthreshold Vm dynamics in regulating cytosolic Ca2+ rise. Since subthreshold Vm depolarization is intrinsically linked to spiking probability, these results are consistent with the finding that Ca2+ rising phase captured higher spike rate (Fig. 3B, D&F).

### Brief intracranial electrical stimulation evoked Vm depolarization is generally accompanied with cytosolic Ca^2+^ increase in CA1 neurons

Our results thus far support a close coupling between intrinsic cytosolic Ca^2+^ increase and subthreshold Vm depolarization in awake mice. To further explore this coupling, we sought to alter Vm and Ca^2+^ through delivering intracranial electrical stimulation via a pair of electrodes on the opposite sides of the imaging window (Fig. 5A) ^49–51^. We delivered either a single electrical pulse, or a burst of 4 pulses (inter-pulse interval: 15 ms) that lasted for a total of 45.2 ms, around the duration of prolonged Vm depolarization observed in CA1 neurons. Each pulse was biphasic, charge balanced, and 100 µs per phase (200 µs total). Due to the variations across animal preparations, the current amplitude was empirically determined in each animal by incrementally increasing stimulation current during 4-pulse stimulation, until a raw GCaMP8m fluorescence change was observed (Methods). The identified current was then used for all subsequent experiments in the given animal, and none of the current used evoked noticeable behavioral effects.

Each neuron was recorded for 16 trials, with each trial randomly assigned as a single pulse or 4-pulse stimulation trial (Fig. 5B). We noted that stimulation led to notable Vm and Ca^2+^ changes across trials in many neurons, though the evoked responses among neurons were quite heterogeneous (Fig. 5C-K, Fig. S9, Fig. S10E1-H1 and I-L), in line with previous observation using voltage or Ca^2+^ imaging alone ^50,51^. Some neurons exhibited immediate Vm hyperpolarization within ∼15 ms of stimulation onset, followed by delayed rebound excitation (Fig. 5C-G), while others showed depolarization with varying temporal profiles within 250 ms of pulse onset (Fig. 5F-K).

To characterize the evoked effect at the individual neuron level, we examined three time windows, 0-15 ms, 15-50 ms, and 50-250 ms after stimulation onset to capture the diverse response profiles. Specifically, we compared the mean Vm during each time window with the corresponding mean Vm during the pre-stimulation period using Wilcoxon Signed Rank test. 16 of the 17 neurons were significantly modulated during at least one time window by either 1-pulse or 4-pulses (Fig. S9Ci and Di). Interestingly, the response evoked by 4-pulse stimulation seems to be generally less prominent than that by 1-pulse (Fig. S9). Within 15 ms of pulse onset, 9 were modulated, and 8 of them were Vm suppressed (Fig. S9Ai and Bi), suggesting that a single electrical pulse leads to immediate Vm hyperpolarization. During the 15-50 ms and 50-250 ms window, 12 were modulated and most exhibited Vm depolarization. As Ca^2+^ imaging was performed at 20 Hz, we did not have the temporal resolution to detect fast Ca^2+^ changes within 50 ms of pulse onset. During the 50-250 ms after pulse onset, neurons with Vm depolarization generally showed an increase in Ca^2+^ (Fig. S9A-B). Additionally, in a subset of neurons, we compared the effect of brief intracranial electrical stimulation across awake, isoflurane anaesthetized, and post-anesthesia awake states. We found that stimulation, either with 1- or 4-pulses, was less effective in evoking Vm and Ca2+ changes during anesthesia (Fig. S10B-D, Fig. S10E-H), consistent with the general idea that isoflurane anesthesia reduces neuronal excitability.

To further evaluate the relationship of evoked Vm and Ca^2+^, we identified the peak amplitude of evoked response during 0-250 ms after stimulation onset. Despite the heterogeneity across neurons, the evoked peak Vm and Ca^2+^ amplitudes exhibited a strong linear relationship for both 1-pulse and 4-pulse stimulation (Fig. 5L&M). This resembles the pattern observed during spontaneous dynamics (Fig. 4K&M), suggesting that brief electrical stimulation likely engages cellular mechanisms that couple Vm and Ca^2+^.

### Prolonged intracranial electrical stimulation leads to diverse Vm and cytosolic Ca^2+^ changes and decouples Vm and Ca^2+^ in many CA1 neurons

To evoke more diverse and longer lasting changes, we additionally tested 0.7 s long trains of electrical stimulation pulses delivered at 40Hz, 140 Hz and 1 kHz that have been shown to evoke heterogeneous CA1 cellular responses^49–51^. Like the brief stimulation experiment, each pulse was biphasic, charge balanced, and 100 µs per phase (Fig. 6A). The current amplitude was again empirically determined in each animal by incrementally increasing stimulation current at 40 Hz until Vm entrainment was detected (Methods), and the identified amplitude was then used for all stimulation frequencies in the given mouse. None of the stimulation conditions produced noticeable behavioral effects.

**Figure 6.**
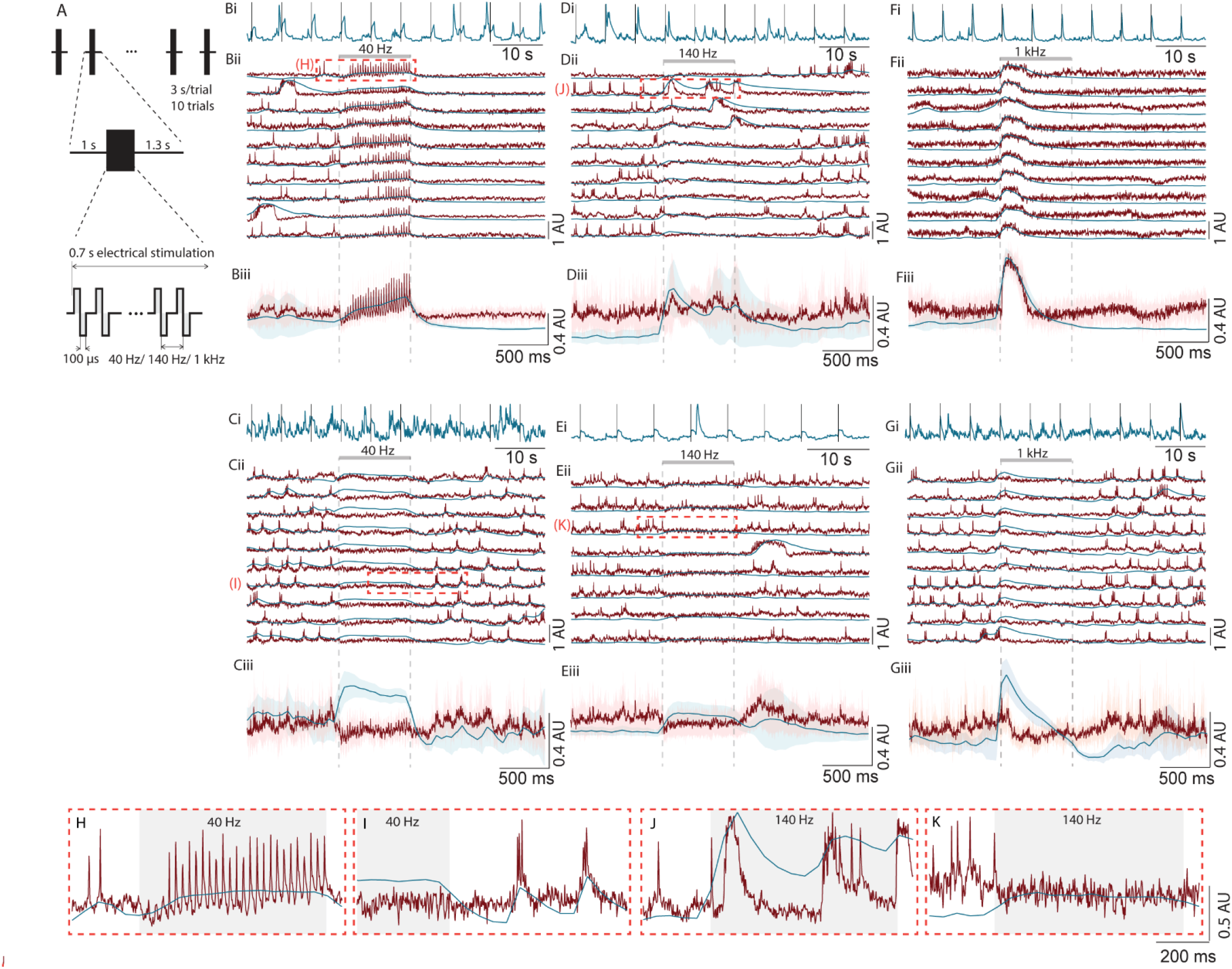
Prolonged electrical stimulation evoked heterogeneous changes in Vm and Ca^2+^ across CA1 neurons in awake mice. (**A**) Prolonged electrical stimulation protocols. Each trial consisted of 1 s baseline, 0.7 s stimulation, and 1.3 s post-stimulation. During stimulation, electrical pulses were delivered at 40 Hz, 140 Hz or 1 kHz, with each pulse being 100 µs/phase. (**B-G**) Example neurons showing significant stimulation-evoked Vm depolarization (B, D, F) and Vm hyperpolarization (C, E, G) upon stimulation at 40 Hz (B, C), 140 Hz (D, E) and 1 kHz (F, G). (**Bi-Gi**) Continuous 1-minute long Ca^2+^ trace of each neuron, with the gray lines indicating stimulation onset. (**Bii-Gii**) Vm (red) and Ca^2+^ (blue) traces across individual trials of each neuron. Stimulation period is highlighted by the gray bar on the top. (**Biii-Giii**) Mean evoked response across trials shown in (Bii-Gii). Shaded areas are standard deviation. Stimulation onsets and offsets are indicated by dashed gray vertical lines. (**H-K**) Zoom-in view of the periods highlighted by red dotted boxes in (Bii-Eii). Gray shadings indicate the stimulation periods. 1 AU=100% dynamic range.

We found that across repeated trials electrical stimulation evoked consistent changes in Vm and Ca^2+^ in many neurons, though the exact responses varied between neurons (Fig. 6B-G). Most neurons (90%) showed robust Vm entrainment during 40 Hz stimulation, but not during 140 Hz stimulation (10% entrained, Fig. 6B-E, Fig. S10 and Table S2), consistent with our previous study using SomArchon alone ^50^. The slow sampling rate of GCaMP imaging at 20 Hz was insufficient for Ca^2+^ entrainment analysis. Thus, we focused on comparing the evoked Vm and Ca^2+^ changes over the entire stimulation period. We observed that stimulation-evoked Vm depolarization was generally accompanied by Ca^2+^ rise in many neurons (Fig. 6B, D&F), with some showing sustained elevation throughout the stimulation period (Fig. 6B) and others showing transient changes following stimulation onset (Fig. 6F). Interestingly, in some neurons, stimulation led to profound Vm hyperpolarization, which however was accompanied by prolonged increase in Ca^2+^ (Fig. 6C, E&G).

To assess the relationship between stimulation-evoked Vm and Ca^2+^ changes, we first determined whether the Vm of a neuron responded to stimulation. We compared the mean Vm during stimulation on each trial versus the corresponding periods before stimulation onset using Wilcoxon Signed Rank test. Neurons showed significant Vm changes across trials were deemed as modulated, either activated or suppressed. We found that all three frequencies modulated a large fraction of neurons, with 12-35% exhibiting significant Vm depolarization (Vm-activated), 42-45% showing Vm hyperpolarization (Vm-suppressed) (Fig. 7Ai and Table S2).

**Figure 7.**
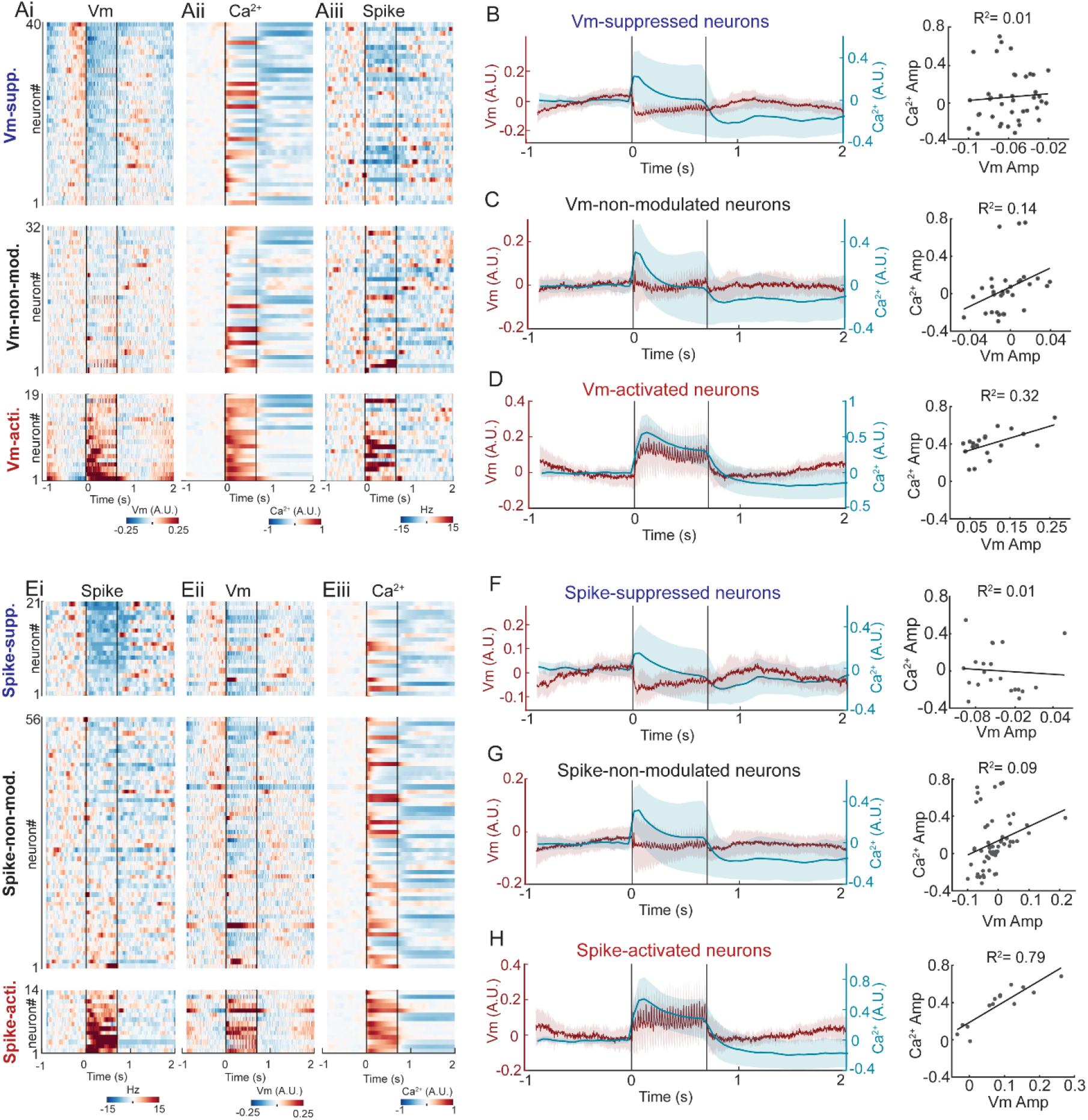
Vm and Ca^2+^ coupling in many neurons was disrupted by prolonged intracranial electrical stimulation. (**Ai**) Heatmaps of stimulation-evoked Vm changes across neurons, aligned to stimulation onset. Each row corresponds to a neuron. Redder color corresponds to stronger Vm depolarization and bluer color corresponds to greater Vm hyperpolarization. Neurons were grouped as Vm-suppressed (top), Vm-non-modulated (middle), or Vm-activated (bottom). Within each group, neurons were sorted by mean Vm changes during the stimulation period in ascending order. (**Aii, Aiii**) Corresponding heatmaps of evoked changes in Ca^2+^ (Aii) and spike rate (Aiii), with the same neuron sorting order as in (Ai). (**B**) Left: mean evoked Vm (red) and Ca^2+^ (blue) across Vm-suppressed neurons. Shaded areas indicate standard deviation. Right: The mean amplitude of evoked Vm versus Ca^2+^ in Vm-suppressed neurons (n=40 neurons, R^2^=0.01). Line indicates linear regression with the R^2^ indicated. (**C**) Similar to (B), but for Vm-non-modulated neurons (n=32 neurons, R^2^=0.14). (**D**) Similar to (B), but for Vm-activated neurons (n=19 neurons, R^2^=0.32). (**Ei**) Heatmaps of stimulation-evoked firing rate changes across neurons, grouped as Spiking-suppressed (top), Spiking-non-modulated (middle), or Spiking-activated (bottom). Neurons were sorted by evoked spike rate changes during the stimulation period in ascending order. Note, the plot is a resorted version of the heatmap shown in Aiii. (**Eii, Eiii**) Corresponding heatmaps of evoked changes in Vm (Eii) and Ca^2+^ (Eiii), with the same sorting order as in (Ei). **(F-H)** Similar to (B-D), but for Spike-suppressed neurons (F, R^2^=0.01), Spike-non-modulated neurons (G, R^2^=0.09), and Spike-activated neurons (H, R^2^=0.79). 1 A.U.=100% dynamic range.

We then grouped all neurons based on their evoked Vm responses regardless of stimulation frequencies. Among Vm-activated neurons, most showed concomitant increase in cytosolic Ca^2+^ (Fig. 7Aii), and the mean amplitude of evoked Vm and Ca^2+^ exhibited moderate correlations (R^2^=0.32, Fig. 7D), largely consistent with that observed during spontaneous Vm depolarization (Fig. 4). However, across Vm-suppressed neurons, many showed a pronounced Ca^2+^ elevation, especially during 40 Hz stimulation (Fig. 7Ai, Aii and Table S2). Among the Vm-suppressed population, the mean amplitudes of evoked Vm and Ca^2+^ showed no correlation (R^2^=0.01), indicating a decoupling between Vm and cytosolic Ca^2+^ (Fig. 7B). In the Vm-non-modulated neurons, both transient and sustained Ca^2+^ elevations were observed (Fig. 7Ai, Aii & C), and the mean amplitudes of evoked Vm and Ca^2+^ showed little correlation (R^2^=0.14).

Further evaluation of stimulation-evoked firing rate change revealed that 48-50% neurons were modulated by 40 Hz or 140 Hz stimulation, showing balanced proportions of activation and suppression (Table S2). In contrast, only 26.2% of neurons were spike-modulated by 1 kHz stimulation, dominated by suppression (19.0% of the 26.2%). Among spike-activated neurons, the mean amplitude of evoked Vm also exhibited a high correlation with the corresponding mean amplitude of Ca^2+^ (R^2^=0.79, Fig. 7H). However, among spike-suppressed neurons and spike-non-modulated neurons, Vm and Ca^2+^ showed no correlation (Spike-suppressed: R^2^=0.01; Spike-non-modulated: R^2^=0.09) (Fig. 7F&G).

Similar analysis categorizing evoked responses based on Ca^2+^ dynamics revealed that a large fraction of neurons exhibited increased Ca^2+^ levels (45-59% for 40 Hz and 140 Hz, 23% for 1 kHz, Table S2), though the evoked Ca^2+^ amplitude varied substantially across neurons (Fig. 6Bi-Gi). Interestingly, we also detected a substantial proportion of neurons showing robust decrease in Ca^2+^, mainly during 1 kHz electrical stimulation (7% for 40 Hz, 0% for 140 Hz, 44% for 1 kHz, Fig. S11Ai and Table S2). Since prolonged reduction in cytosolic Ca^2+^ is not common during spontaneous activities, the evoked Ca^2+^ suppression suggests that electrical stimulation engages cellular signaling that is not typically active during regular physiological processes. The fact that 1 kHz stimulation led to robust Ca^2+^-suppression also highlights distinct cellular mechanisms engaged by 1 kHz stimulation compared to 40 Hz and 140 Hz. Ca^2+^-activated neurons exhibited diverse changes in Vm, and the mean amplitude of evoked Ca^2+^ showed no correlation with the corresponding Vm (R^2^=0.01) (Fig. S11Ai, Aii&D). Similarly, Ca^2+^-suppressed neurons, dominated by 1 kHz stimulation condition, also displayed divergent Vm changes with no correlation (R^2^=0.02, Fig. S11B).

Overall, these results demonstrate that Vm depolarization evoked by electrical stimulation is closely accompanied by cytosolic Ca^2+^ rise, similar to that observed under spontaneous conditions. In contrast, stimulation-evoked Vm hyperpolarization is decoupled from cytosolic Ca^2+^. These results support the notion that the cellular mechanisms underlying Vm depolarization and cytosolic Ca^2+^ rise are intrinsically linked. The non-physiological Vm hyperpolarization during prolonged electrical stimulation may engage membrane ionic currents or intracellular Ca^2+^ resources that disrupt the coupling of Vm and cytosolic Ca^2+^ dynamics.

## Discussion

Membrane voltage (Vm) exhibits complex dynamics that regulate spike timing and cellular signaling during behavior ^3,52^. However, it is unclear how Vm in the awake mammalian brain engages Ca^2+^-dependent signaling pathways known to be critical for cellular function and plasticity. Accordingly, we measured Vm and Ca^2+^ simultaneously in the same neurons using the color-compatible, genetically encoded voltage sensor SomArchon and Ca^2+^ sensor GCaMP. We found that in the CA1 and striatum of awake mice, only ∼28% of spikes were accompanied by Ca^2+^ events, in sharp contrast to the ∼70% observed in rat and human iPSC-derived neurons that exhibit little Vm fluctuations. Furthermore, in the awake brain, slow subthreshold Vm depolarization is tightly coupled to Ca^2+^ increases, particularly during complex spikes (CS), suggesting that slow Vm dynamics robustly engages cytosolic Ca^2+^ and tracks changes in firing rate. While plateau potential has been implied to be accompanied by dendritic Ca^2+^ changes, the supporting evidence has been indirect, either from separated Ca^2+^ and Vm recordings in awake mice ^20,53,54^ or simultaneous recordings in anesthetized brains using current injection to produce artificial Vm depolarization ^24,55^. By deploying a novel AAV viral construct to measure Vm and Ca^2+^ in the same neurons in awake mice, our study provides direct experimental evidence of a tight coupling between prolonged subthreshold Vm depolarization and large Ca^2+^ rise in awake brains. Interestingly, Ca^2+^ increase was also more evident after SS-w-ADP than after SS-wo-ADP, further supporting the coupling of Ca^2+^ with slow Vm dynamics. Thus, slow subthreshold Vm depolarization is faithfully translated into Ca^2+^-mediated cellular changes, and different spiking modes produce distinct cellular consequences via selective regulation of cytosolic Ca^2+^. Finally, Vm and Ca^2+^ coupling was disrupted in a sizable fraction of CA1 neurons by prolonged intracranial electrical stimulation, broadly used in clinical neuromodulation, underscoring the potential to differentially regulate neuronal output versus cellular signaling.

While spiking is critical for neural circuit computation, subthreshold Vm dynamics determine the timing of spiking. Both spiking and subthreshold dynamics are dictated by the biophysical properties of a neuron and shaped by cellular plasticity that typically involves Ca^2+^-dependent signaling pathway ^6,56,57^. Subthreshold Vm depolarization, once reaching threshold for action potential initiation, leads to spiking, which in turn modulates diverse voltage-gated Na^+ 31^, K^+ 58^, and Ca^2+ 59,60^ channels to further influence Vm and subsequent spiking. For example, action potential width increases and inter-spike-interval lengthens in later spikes within a CS ^3,26^, which have been related to spike-induced inactivation of Na^+^ channels ^32,61^. With SomArchon voltage imaging, we were able to investigate the relationship between subthreshold Vm and Ca^2+^ in awake mice. This has not been feasible with traditional electrophysiology techniques that only measure action potentials extracellularly. In addition to CS that were accompanied by prolonged ADP and large Ca^2+^ increase, we also detected a sizable fraction of SS-w-ADP with small but persisted Ca^2+^ increase (Fig. 4R-S), in contrast to SS-wo-ADP that was accompanied by minimal Ca^2+^ fluctuation (Fig. 4T). To detect ADP following each spike, we compared the mean Vm amplitude during 5-30ms after spike versus before spike. Using more strict criteria to identify ADPs could reveal additional cytosolic Ca2+ patterns that may have been missed here. These results demonstrate differential coding mechanisms among isolated spikes, beyond previously described differences between spike bursts and single spikes. It is conceivable that SS-w-ADP represents an intermediate between CS and SS-wo-ADP, and these different spiking patterns correspond to a continuum of Vm dynamical states. It is possible that the transition between these spiking modes depends on the amplitude and duration of the ADP, which is shaped by intracellular Ca^2+^ and other signaling pathways. Future analyses of different types of SSs based on their association with ADPs or Ca^2+^ will uncover their specific circuit and behavioral impacts.

In addition to the systematic analysis of the relationship between spiking and cytosolic Ca^2+^ from 4-21 neurons across four mouse lines ^12^, several other studies also performed simultaneous electrophysiology and Ca^2+^ imaging in a small number of neurons in anesthetized mice ^8,62,63^. Example traces from these studies follow similar patterns, where spike bursts are generally accompanied by larger Ca^2+^ events, whereas isolated single spikes are followed by smaller Ca^2+^ changes. It is highly plausible that the magnitude of cytosolic Ca^2+^ concentration change during spiking follows a continuum, and thus the 0-1 spike conversion of Ca^2+^ events have intrinsic limitation, especially given variations among recordings and experimental conditions where GCaMP expression and other factors impact fluorescence detections.

CSs are traditionally classified as bursts of spikes superimposed on prolonged Vm depolarization ^24^. However, patch clamp studies have shown that some CSs may contain only one spike during the prolonged subthreshold depolarization, while others may have a few spikes spaced by tens of milliseconds in between ^64^. Our recordings contain diverse CS patterns, and they could co-exist in one neuron (Fig. 4B1&B2 and C1&C2). While Vm depolarization and spike probability are intricately linked, their generation involve distinct biophysical mechanisms. For example, blocking spiking with Na^+^ channel blockers failed to eliminate prolonged Vm depolarization ^24^. Furthermore, spiking and prolonged Vm depolarization (plateau potentials) in CA1 place cells are shaped by behavioral experience ^20^. Milicevic *et al* found that subthreshold Vm depolarizations (>10 mV and >200 ms) could generate conspicuous Ca^2+^ events *in vitro* ^38^. We directly probed the relationship between Ca^2+^ and slow Vm in awake mice (Fig. 4I), and detected a close correlation between Ca^2+^ rise and slow Vm depolarization. Our observation in the awake mouse brain is consistent with previous in vitro observation that subthreshold Vm depolarizations led to large Ca^2+^ events ^38^. Since slow Vm depolarization relates to intrinsic membrane excitability ^17,58^, our results provide, to our knowledge, the first direct *in vivo* experimental evidence that somatic Ca^2+^ is better at tracking neuronal excitability over a slower time scale than spike timing at the millisecond time scale. Furthermore, Owen *et al.*^43^ reported that inhibition of striatal fast-spiking interneurons led to large Ca^2+^ changes in striatal medium spiny neurons that cannot be easily explained by spiking changes. It is possible that prolonged Vm depolarization in medium spiny neurons (Fig. S5F-I), traditionally referred to as “up states” in electrophysiology recordings, may underlie the robust increase in cytosolic Ca^2+^ upon inhibition of local interneurons reported by Owen *et al*.

As we imaged neuronal soma, the Ca^2+^ rise after Vm depolarization likely activates somatic Ca^2+^-dependent calmodulin pathways that are known to be important for cellular plasticity ^1,57^. Slow somatic Vm depolarization relies on dendritic Ca^2+^ channels ^19,20^, and Ca^2+^ dynamics has been shown to be highly correlated between the soma and dendritic compartments ^65^. Thus, the observed somatic Ca^2+^ elevation during Vm depolarization could be accompanied by dendritic Ca^2+^ increases, which in parallel regulate synaptic plasticity in dendrites. As the delay between Ca^2+^ rise after Vm depolarization is on the order of tens of milliseconds (Fig. 4J&L), our results suggest that synaptic plasticity can be precisely regulated by Vm.

Cytosolic Ca^2+^ elevation can be attributed to direct Ca^2+^ influx through the plasma membrane via voltage-gated Ca^2+^ channels or indirect Ca^2+^ release from endoplasmic reticulum ^57^, mitochondria ^66^, and other intracellular Ca^2+^ stores ^67^. In the cortex, hippocampus, and striatum, neurons generally express all common types of voltage-gated Ca^2+^ channels ^14,68,69^. The high-voltage, spiking-activated L-type Ca^2+^ channels have been shown to contribute to Ca^2+^ influx during ADPs and place field formation^6,57,70^. Similarly, the low-voltage-gated T-type Ca^2+^ channels, activated by subthreshold Vm depolarization and inactivated by spiking, regulate spike initiation and bursting within CS ^13^. Intriguingly, we noted a very small Ca^2+^ rise preceding SS-wo-ADP (Fig. 4H and T, Fig. S5E and Fig. S6E), which may reflect T-type Ca^2+^ channel activation. T-type Ca^2+^ channels additionally could contribute to the Ca^2+^ increase during spike bursting in cortical PV cells following prolonged Vm hyperpolarization (Fig. 2R-S) ^28,71^. In addition to direct Vm-dependent Ca^2+^ influx through the plasma membrane, dendritic Ca^2+^ increases mediated by NMDA receptors could also contribute to slow Vm depolarization during CSs ^72–74^. Cytosolic Ca2+ and subthreshold Vm are intricately linked. Grienberger et al. showed that dendritic NMDA receptors and the voltage-gated Ca^2+^ channels are essential for slow Vm depolarization that drives CS ^24^, distinct from the Ca^2+^ influx observed after spiking. Recently, Gonzalez *et al.* found that CS propagates further into dendrites than SS, suggested that CS and SS engage different intracellular signal pathways ^75^. Finally, the Ca^2+^-mediated Ca^2+^ release from intracellular stores, such as mitochondria and endoplasmic reticulum ^57^, could further amplify cytosolic Ca^2+^ rise, extending Ca^2+^ peak delays after Vm depolarization.

The intracranial electrical stimulation pulses used were charge-balanced biphasic square waves, with a width of 100 µs/phase, which are clinical standard parameters. Using these parameters, we found that prolonged stimulation evoked both Vm depolarization and hyperpolarization (Fig. 6 and Fig. 7Ai). While evoked Vm depolarization was generally accompanied by Ca^2+^ rise, Vm hyperpolarization frequently decoupled from the expected Ca^2+^ reduction, manifesting instead as an Ca^2+^ increase. Typically, Vm hyperpolarization is driven by inhibitory inputs, and the resulting influx of anions, such as Cl^−^, which would not activate voltage-gated Ca^2+^ channels. Conversely, elevated intracellular Ca^2+^ can itself drive Vm hyperpolarization by activating Ca^2+^-dependent K^+^ channels ^76^. The observed decoupling suggested a divergence in the cellular mechanisms governing Vm and Ca^2+^ homeostasis during the prolonged electrical stimulation. For example, cytosolic Ca^2+^ clearance via Ca^2+^-ATPases ^15,66^, which pump cytosolic Ca^2+^ out of the cell or into the endoplasmic reticulum and mitochondria, may be attenuated by the electrical stimulation pulse trains. Furthermore, electrical stimulation likely recruits neuromodulatory systems, triggering the release of ligands that engage GPCR signaling cascades ^77,78^. These pathways can trigger Ca^2+^ release into the cytosol from intracellular stores via the IP_3_ receptor pathway ^79^ or facilitate Ca^2+^ influx through non-selective cation channels ^18^, both of which operate independent of the instantaneous Vm state.

Transient GCaMP fluorescence increase has been anecdotally used as a surrogate for spiking. We found that Ca^2+^ event rising phase only captured ∼28% of spikes in the awake brain, consistent with the previous study by Huang *et al.* ^12^ showing that GCaMP6 captured <10% of spikes and ∼20-30% of two-spike bursts using extracellular recordings and two-photon Ca^2+^ imaging in anesthetized mice. *In vitro*, we found that Ca^2+^ events captured ∼70% of action potentials. This difference may be attributed to the prominent subthreshold Vm dynamics in the brains of awake mammals, which modulates the interplay between Vm and Ca^2+^, leading to more complex relationship than the tightly coupled spike-Ca^2+^ dynamics observed *in vitro* with limited subthreshold dynamics. However, we did observe occasional prolonged subthreshold Vm depolarization in cultured human neurons (Fig. 1O), which might be attributed to the >4 weeks culture time that allows for more substantial synapse formation than rat neurons. Finally, by comparing the evoked Vm and Ca^2+^ responses to brief electrical stimulation in the same neurons across awake, anaesthetized, and post-anesthesia awake states, we found isoflurane anesthesia substantially reduced neuronal excitability (Fig. S9), highlighting the importance of model choices.

There are many other differences between *in vivo* and *in vitro* conditions, and the strong fluorescence background *in vivo* may also limit GCaMP’s ability to detect small Ca^2+^ increases associated with individual spikes. Milicevic *et al.* ^80^ found that 500 Hz sampling rate improved the detection of Ca^2+^ rising phase compared to 14 Hz sampling rate, captured smaller amplitude Ca^2+^ transients and detected more Ca^2+^ events. However, a higher sampling rate of 99 Hz did not improve spiking event detection by Ca^2+^ events using the Ca^2+^ indicator jRGECO1a ^81^. Consistent with Milicevic *et al*., we also noted a higher Ca^2+^ event rate under the higher 50 Hz sampling rate condition (Table S1). However, higher LED power (∼150 mW/mm^2^) was needed to image GCaMP at 50 Hz sampling rate than 20 Hz (∼40 mW/mm^2^), leading to significantly greater photobleaching (Fig. S4C-H), which limited recording time. Besides the influences by imaging techniques, the non-linearity, binding affinity, temporal resolution and sensitivity of fluorescence indicators limits the ability to probe finer relationship between Vm and Ca^2+^.

## Materials and Methods

### Molecular cloning and AAV production

Archon1 (GenBank ID MG250280.1), jGCaMP7f (GenBank ID MK749392.1) and jGCaMP8m (GenBank ID OK646319.1) were assembled into the pAAV2 viral vector backbones to obtain the following constructs: AAV-Syn-SomArchon-P2A-GCaMP7f, AAV-Syn-SomArchon-P2A-GCaMP8m, and AAV-CAG-FLEX-SomAchon-P2A-GCaMP8m. Molecular cloning was performed by Epoch Life Science, Inc. AAV2.9 viral particles were packaged by Charles River Laboratories (Previously, Vigene Biosciences) with a titer of 3.7×10^12^ GC/mL for AAV9-Syn-SomArchon-P2A-GCaMP7f, 1.8×10^13^ GC/mL for AAV9-Syn-SomArchon-P2A-GCaMP8m, and 2.7×10^13^ GC/mL for AAV9-CAG-FLEX-SomAchon-P2A-GCaMP8m.

### Custom wide-field microscope for simultaneous SomArchon and GCaMP imaging

Simultaneous SomArchon and GCaMP imaging was performed using a custom wide-field microscope configured with dual optical pathways. Excitation was provided by a 470 nm LED (Thorlabs, M470L4) for GCaMP and a 637 nm laser (Ushio America, Necsel Red 63×) for SomArchon (Fig. 1A and Fig. S1). For GCaMP excitation, the 470 nm LED light was conditioned through an aspherical condenser (L4 in Fig. S1, f = 16 mm, Thorlabs ACL25416U). For SomArchon excitation, the 637 nm laser was directed though a high-NA achromatic collimator (L1 in Fig. S1, Thorlabs F950FC-A), followed by an achromatic doublet lens (L2 in Fig. S1, f=200 mm, Thorlabs AC254-200-A) and an Iris diaphragm for mechanical regulation of laser intensity. These excitation beamss were coaxially combined via a 505 nm long-pass dichromatic mirror (DM1 in Fig. S1, Thorlabs, DMLP505R), which reflected the 470 nm light and transmitted the 637 nm light.

The combined two excitation beams passed through an achromatic doublet lens (L3 in Fig. S1, f=200 mm, Thorlabs AC254-200-A), filtered by a fluorescence excitation filter (F1 in Fig. S1, FF01-390/482/532/640-25), reflected by a quadband dichromatic mirror (DM2 in Fig. S1, Semrock, Di03-R405/488/532/635), and focused onto the sample though a 40X objective (Nikon, CFI Apo NIR 40×/0.8 NA W).

The fluorescence from the sample was collected by the objective, passed through DM2, and filtered by a multi-band emission filter (F2 in Fig. S1, Semrock FF01-446/510/581/703-25). For simultaneous dual-color detection, a multi-band dichromatic mirror (DM3 in Fig. S1, Chroma Technology Corp., ZT405/514/635rpc) split the fluorescence, reflecting GCaMP fluorescence and transmitting SomArchon fluorescence into separate detection paths.

In the reflected path (GCaMP), the fluorescence was directed through an achromatic doublet lens (L5 in Fig. S1, f=200 mm, Thorlabs AC254-200-A) and a 525±22.5 nm bandpass filter (F3 in Fig. S1, Semrock, FF01-525/45), and then was recorded via an sCMOS camera (Hamamatsu Photonics, C13440-20CU). In the transmitted path (SomArchon), the fluorescence passed through an achromatic doublet lens (L6 in Fig. S1, f=200 mm, Thorlabs AC254-200-A) and a 700±37.5 nm bandpass filter (F4 in Fig. S1, Chroma Technology Corp., ET700/75m), and then was recorded by a second sCMOS camera (Hamamatsu Photonics, C15440-20UP). The intensity of 637 nm laser excitation on the imaging field was ∼6 W/mm^2^, and the intensity of the blue LED excitation was ∼40 mW/mm^2^.

To minimize SomArchon photobleaching, a mechanical shutter (Thorlabs, SH1) and an adjustable iris (Thorlabs, SM1D12D) was positioned in the 637 nm excitation path to control the timing and the excitation area (∼50 μm in diameter). We confirmed the separation of the two fluorescence light paths using non-labelled brain tissue and a piece of Kimwipe. Due to non-specific autofluorescence, some very weak and static fluorescence was detected on the GCaMP light path (∼0.3% of the baseline intensity) during 637 nm laser illumination, and on the SomArchon light path (∼2% of the baseline intensity) during 470 nm LED illumination.

### Rat primary neuron culture preparation

Rat hippocampal neuron cultures were prepared from 18-day rat embryos as previously described ^82,83^. Briefly, isolated hippocampal neurons were plated on poly-L-lysine (0.1 mg/mL, Sigma-Aldrich, P2636-100MG) coated coverslips (Electron Microscopy Sciences, 72296-08, #1.5, diameter: 8 mm) at a density of 100 k/well on 24-well plates. The plating media contained 10% FBS, 5% heat-inactivated horse serum and 260 µM cystine in DMEM with L-glutamine. One day post plating, the plating media were replaced with Neurobasal media containing 1% 100× Penicillin-Streptomycin, 2 mM L-glutamine, 1% donor horse serum and 2% SM1 neuronal supplement (STEMCELL Technologies, 05711). Cells were maintained at 37°C with 5% CO_2_. Neurobasal media was exchanged twice a week. Cells were transduced with AAV9-Syn-SomArchon-P2A-GCaMP7f (∼7×10^9^ GC/well) 2 days after plating.

### Human iPSC-derived neuron culture preparation

Human neuroepithelial stem cells (hNESCs) were derived from human iPSC (healthy, female, age at sampling 81 years). The lines have been described first time by Reinhardt *et al.* ^84^ and cultured according to the procedures described there. Human iPSC-derived neurons were generated from neuroepithelial stem cells as described in Monzel *et al.* ^85^. Briefly, neuroepithelial stem cells were plated on Matrigel (Matrigel hESC-Qualified Matrix, Corning, 354277; Knockout™ DMEM, Thermo Fisher, 10829018) coated 6-well plates at a density of 500 k/well, and maintained in the maintenance media containing an equal mix of DMEM-F12 (Thermo Fisher, 21331020) and Neurobasal medium (Thermo Fisher, 21103049) supplemented with 1% GlutaMAX™ Supplement (Thermo Fisher, 35050061), 1% Gibco™ Penicillin-Streptomycin (10,000 U/mL, Thermo Fisher, 15140122), 1% B27 Supplement 50X (minus vitamin A, Thermo Fisher, 12587010), 0.5% N2 Supplement 100X (Thermo Fisher, 17502048), 150 µM L-ascorbic Acid (Sigma-Aldrich, A5960-25G), 3 µM CHIR 99021 (Axon Med Chem, 1386), and 0.75 µM purmorphamine (Sigma-Aldrich, SML0868). The maintenance media was refreshed every three days after plating. Seven days after plating, the media was replaced with the differentiation media (Day 0 of differentiation), containing N2B27 media supplemented with 10 ng/mL hGDNF (PeproTech, 450-10-10µg), 10 ng/mL hBDNF (PeproTech, 450-02-10µg), 1 ng/mL recombinant human TGF-β3 (PeproTech, 100-36E-10µg), 500 µM N6,2’-O-Dibutyryladenosine 3′,5′-cyclic monophosphate sodium salt (db-cAMP, Millipore Sigma, D0627-250MG), 200 µM ascorbic acid, and 1 µM purmorphamine. Four days later, cell cultures were digested using StemPro™ Accutase™ Cell Dissociation Reagent (Thermo Fisher, A1110501), suspended, and then replated in differentiation media on Matrigel-coated coverslips in 24-well plates at a density of 80-120 k/well. Differentiation media were replaced twice a week. From Day 7 and onward, differentiation media without purmorphamine was used. Cells were transduced with AAV9-Syn-SomArchon-P2A-GCaMP7f on Day 7-10 post differentiation at ∼7×10^9^ GC/well.

### Animal preparation

All animal experiments were performed in accordance with the National Institute of Health Guide for Laboratory Animals and approved by the Boston University Institutional Animal Care and Use Committee (IACUC). 23 adult C57BL/6 mice (Charles River Laboratories Inc.) were used (10 female and 8 male mice for CA1 experiments, and 3 female and 2 male mice for dorsal striatum experiments). 3 PV-cre mice (2 male and 1 female, Jackson Laboratory, JAX stock #017320, B6.129P2-Pvalb^tm1(cre)Arbr^/J) were used for the V1 experiments. All mice were 8-22 weeks of age at the start of the study.

Custom imaging windows for CA1 and striatum experiments were constructed by affixing a circular glass coverslip (Warner Instruments Inc., 64-0726 (CS-3R0), #0, OD: 3 mm) to the bottom of a stainless-steel cannula (AmazonSupply, B004TUE45E, OD: 3.17 mm, ID: 2.36 mm, height: 1.75 mm) using UV-curable glue (Norland Products Inc., Norland Optical Adhesive 60, P/N 6001). An 8 mm guide cannula (PlasticsOne Inc., C135GS-4/SPC, 26 G) was then coupled to the side of the imaging window using solder (RadioShack, 6400013). We additionally affixed two polyimide insulated stainless steel electrode wires (PlasticsOne Inc., 005SW/30S WIRE 37365 S/S .005” DIA. OPTION-01) along the infusion cannula, using Epoxy (Thorlabs, G14250), with the electrode tips terminating underneath the center bottom of the imaging window. Approximately 0.5 mm of the electrode tip was exposed by removing the wire insulation to reduce electrode impedance for electrical stimulation. The two electrode tips were ∼3 mm apart.

Under general anesthesia with isoflurane (∼1-3%), we surgically implanted the imaging window over the hippocampal CA1 (centered at the stereotactic coordinates: M/L: 1.7 mm, A/P: −2 mm, D/V: −1.4 mm), after gently removing the overlaying cortex, the corpus callosum, and thinning the alveus layer. During the same surgery, a metal pin (DigiKey, ED85100-ND) soldered to a screw (J.I. Morris Company, F000CE094) was placed over the left cerebellum to serve as the ground for impedance measurements for the stimulation electrodes. The imaging window, the ground pin, and a custom headbar were affixed to the skull posterior to the imaging window using bone cement (Parkell Products Inc., C&B Metabond) and dental cement (Stoelting, 51459). Similar surgical protocols were used to implant the imaging window over the striatum (centered at stereotactic coordinates: M/L:1.8 mm, A/P: 0.5 mm, D/V: 2.2 mm), after carefully thinning the corpus callosum to expose the dorsal striatum.

Ten to fourteen days post-surgery, we infused a total of 3-5×10^10^ AAV viral particles per mouse, corresponding to 2-6 µL depending on the titer of the virus, through the implanted guide cannula using an infuser (PlasticsOne Inc., C315IS-4/SPC, 33 G) coupled to a 10 μL syringe (World Precision Instruments LLC, NANOFIL). Each infusion lasted for 15min, and the exact infusion speed was adjusted by the total volume infused (0.1-0.4 µL/min) and controlled via an infusion pump (World Precision Instruments LLC, UltraMicroPump3-4). At the completion of the infusion, the infuser was left in place for 10 min before removal.

To image PV cells in V1, we first created a craniotomy of ∼3 mm in diameter (centered at stereotactic coordinates: M/L: 2.3 mm, A/P: −3.6 mm), and then injected AAV9-CAG-FLEX-SomArchon-P2A-GCaMP8m into three sites within the craniotomy (500 nL per spot) via a 10 μL syringe (World Precision Instruments LLC, NANOFIL) controlled by a pump (injection speed: 80 nL/min, World Precision Instruments LLC, UltraMicroPump3-4). After viral injection, we affixed a circular glass coverslip (Warner Instruments Inc., 64-0726 (CS-3R0), #0, OD: 3 mm) over the craniotomy using bone cement (Parkell Products Inc., C&B Metabond) and dental cement (Stoelting, 51459). Other surgical and handling procedures were as detailed above for hippocampal and striatal experiments.

### Two-color imaging in awake mice

Imaging in awake mice was performed at least 2 weeks after viral infusion to allow for sufficient SomArchon and GCaMP expression. During each imaging session, mice were head-fixed and voluntarily locomoting on a treadmill made of an air floating ball as detailed previously ^86^. We first identified the area of interest using GCaMP fluorescence, and then collected GCaMP and SomArchon fluorescence simultaneously using two sCMOS cameras. Specifically, each of the two sCMOS cameras was controlled by a separate computer running HCImage Live (Hamamatsu Photonics K.K.). For each field of view (FOV), GCaMP fluorescence was continuously recorded for 1-2 minutes, while SomArchon fluorescence was recorded intermittently for half of the time, resulting in 5-10 trials, with each trial lasting for 3-6 seconds every 6-12 seconds.

SomArchon fluorescence images were collected using HCImage, triggered via a TTL pulse generated by a DAQ (National Instruments, BNC-2110 and PCIe-6323) controlled by MATLAB. Once triggered, HCImage Live collected 3 or 6 seconds of data (1500 or 3000 frames) with an exposure time set at 2 ms, resulting in an actual frame capture every 2.012 ms, as determined using the DAQ at 250 kHz acquisition rate. At 500 frames per second, the amplitude of spikes is substantially smaller than that achieved by conventional patch clamp recording. HCImage was set to capture 144×144 or 144×288 pixels with 2×2 binning, corresponding to 46×46 µm^2^ or 46×92 µm^2^. To minimize SomArchon photobleaching, another set of TTL pulses were used to control the shutter in front of the 637 nm laser to open at 0.5 s before the start of image captures and to close at 0.2 s after the end of image captures.

GCaMP fluorescence images were collected using HCImage at 128×128 or 256×256 pixels with 2×2 binning, corresponding to 41×41 µm^2^ or 82×82 µm^2^. Start of GCaMP imaging was triggered by the first TTL pulse generated by the SomArchon camera when SomArchon imaging was first initiated as described above. Once the GCaMP camera was triggered, HCImage Live continuously collected 1 or 2 minutes of data with the exposure time set at 50 ms, resulting in an actual frame capture every 50.008 ms, as determined using the DAQ at 250 kHz acquisition rate. Blue LED for GCaMP imaging was manually switched on and off.

SomArchon and GCaMP image frames were aligned offline after confirming the relative frame timings of the two cameras. Using a DAQ at 250 kHz acquisition rate, we confirmed that the first SomArchon frame captured occurred 1.4 ms after the TTL trigger, and the first GCaMP frame occurred 48.6 ms after the first SomArchon frame. We further verified the timing of the TTL pulses used to trigger SomArchon image frame captures in a subset of recordings using a DAQ at 5kHz acquisition rate.

For higher sampling rate recordings, the exposure time of SomArchon and GCaMP camera was set at 1.5 ms and 20 ms respectively, resulting in an actual frame capture every 1.508 ms and 20.004 ms as determined using the DAQ at 250 kHz acquisition rate. To allow for the higher sampling rate of SomArchon camera, HCImage was set to capture a smaller FOV of 120×144 pixels with 2×2 binning, corresponding to 38.3×46 µm^2^. No FOV size change for GCaMP camera. GCaMP fluorescence was continuously recorded for 64 seconds, while SomArchon fluorescence was recorded intermittently for half of the time, resulting in 16 trials, with each trial lasting for 2 seconds every 4 seconds, with pulse stimulation occurring at the one second after trial starts. All the other setups were the same as described above.

### Two-color imaging in same neurons across awake, anaesthetized, and waking up brain states

First, CA1 neurons were recorded when the mice were awake, with the protocol described above. Second, the mice were left on the ball, head-fixed, and put under general anaesthetization by applying 2% isoflurane through the mask in front of their nose. When the respiration rate lowered down to ∼1 Hz, and the mice showed no response to tail pinch, we located the same neurons that were recorded before the anaesthetization and performed the same recording protocol. After the recording during anaestherization, the mask was removed. In our experiments, the mice started to run on the ball within ∼10-20 min after the isoflurane removal. We relocate the same neurons, and performed recordings during the waking up period (∼40-60 min after the isoflurane removal) with the same protocol. For each neuron, the current amplitude of one or four pulse stimulation remained the same across the three brain states.

### Two-color imaging in cultured neurons

Neuron culture coverslips were transferred to a microscope stage and bathed with Tyrode’s solution (pH=7.4) consisting of 10 mM HEPES (Gibco, 15630-080), 10 mM glucose (Sigma-Aldrich, G8270-100G), 2 mM calcium chloride (Fluka, 21114-1L), 2 mM magnesium chloride (Fluka, 63020-1L), 2.4 mM potassium chloride (Sigma-Aldrich, P9333-500G), and 150 mM sodium chloride (Sigma-Aldrich, S7653-1KG). Fresh Tyrode’s solution was prepared on the day of imaging and filtered with a 0.22 µm Stericup quick release-GP filter system (EMD Millipore, S2GPU05RE). Imaging sessions in rat primary hippocampal neurons were performed 12-14 days post viral transduction. For each FOV, GCaMP fluorescence was continuously recorded for 5 minutes, whereas SomArchon fluorescence was recorded intermittently for half of the time, resulting in 5 trials with each trial lasting for 30 seconds every minute. The HCImage Live software settings were 2 ms per frame (15000 frames) for SomArchon recording and 50 ms per frame (6000 frames) for GCaMP recording. Imaging sessions in human iPSC-derived neurons were performed during Day 20-24 post differentiation. 12-24 hours before imaging, cells were bathed in differentiation media supplemented with all-trans-retinal (5 µM, Sigma-Aldrich, R2500-25MG, stock solution: 50 mM in DMSO). For each FOV, SomArchon and GCaMP fluorescence were collected continuously for 45 seconds. The HCImage Live software settings were 1.5 ms per frame for SomArchon recording and 50 ms per frame for GCaMP recording. The timing of SomArchon and GCaMP image frames were confirmed using DAQ at 250 kHz acquisition rate, and aligned as detailed above for in vivo imaging.

### SomArchon and GCaMP fluorescence image stacks motion correction, ROI selection and trace extraction

All data analysis was performed offline in MATLAB (R2018b, R2021a). SomArchon image stacks were first motion corrected using the NoRMCorre piecewise rigid motion correction algorithm ^87^. As multiple fluorescence imaging trials were obtained for each recording, we first motion corrected all frames within each trial, and then motion corrected the average frames across trials. Image stacks for each trial were first smoothed using a 3-D box filter (MATLAB, imboxfilt3) and a high-pass spatial filter (MATLAB, imfilter) to remove background signals and to create reference points for NoRMCorre image registration. NoRMCorre rows and columns were set to the image stack size, the bin width set to 30, the max shift set to 50, without upsampling, and all other algorithm parameters set to default values. The motion shifts calculated by NoRMCorre were then applied to the all images within a trial using the MATLAB function circshift. Similarly, motion shifts across trials were calculated using average frames computed from the motion corrected individual trials using NoRMCorre (bin width of 1, max shift of 50, and no upsampling). The cross-trial shifts were applied to all image stacks to obtain final motion corrected stacks. GCaMP image stacks were converted to multitiff files first, and then motion corrected as described before ^88^. Briefly, a reference frame was created by averaging the pixel values across all frames. For prolonged electrical stimulation recordings, all pixels in the reference frame were included for motion correction. For all the other recordings, all pixels in a manually drawn rectangular area that covered the neurons were included for motion correction. Motion correction was performed using cross-correlation between each contrast-enhanced and normalized frame and the reference frame.

For SomArchon region of interest (ROI), Individual neurons were manually segmented as a polygonal ROI in MATLAB from the average motion corrected SomArchon image stacks (MATLAB drawpolygon). Raw SomArchon fluorescence traces were extracted by averaging all pixel intensities within each neuron over time. For each SomArchon FOV, a raw background trace was obtained by averaging the pixel intensities of all non-ROI pixels. GCaMP ROIs were selected in imageJ by polygonal selection tools, and imported into MATLAB. The averaged pixel intensity in each ROI was then extracted to obtain raw GCaMP traces. To obtain raw GCaMP background traces, a donut area was created for each GCaMP ROI. The outer and inner boundary of the donut area was 12 and 2 pixels away from the ROI boundary, respectively. The raw GCaMP background trace was obtained by averaging the pixel intensities of the donut area pixels. Each motion corrected SomArchon and GCaMP video was visually inspected. All raw SomArchon and GCaMP traces were manually inspected for signal intensity and recording stability. For *in vivo* analysis under physiological conditions, only neurons with >5 stable recording trials were analyzed. Overall, 31 CA1 neurons from 5 Syn-SomArchon-P2A-GCaMP7f mice, 36 CA1 neurons from 9 Syn-SomArchon-P2A-GCaMP8m mice, and 21 striatal neurons from 5 Syn-SomArchon-P2A-GCaMP8m mice were included in the analysis of spontaneous neuronal activities, 10 CA1 neurons from 3 Syn-SomArchon-P2A-GCaMP8m were included in the analysis of higher sampling rate recordings, 17 CA1 neurons from 3 Syn-SomArchon-P2A-GCaMP8m were included in the analysis of 1 or 4 electrical pulse stimulation effects, and 91 CA1 neurons from 7 Syn-SomArchon-P2A-GCaMP8m mice were included in the analysis of 0.7 s sustained electrical stimulation effects. *In vitro*, trials without spiking were excluded. In total, 11 rat neurons and 16 human iPSC-derived neurons were included in the analysis.

### Vm and Ca^2+^ trace analysis in cultured neurons

We first normalized each SomArchon and GCaMP trace between the maximum (1) and minimum (0) to obtain membrane voltage (Vm) traces and cytosolic calcium (Ca^2+^) traces respectively. Since cultured neurons exhibited little subthreshold Vm fluctuations, only spikes were detected and analyzed. For spike and Ca^2+^ event detection, we first detrended Vm and Ca^2+^ traces by subtracting the baseline calculated as the mean over a 100 data-point moving window. Rapid Vm depolarization corresponding to spikes were identified using the MATLAB findpeaks function (‘MinPeakHeight’=0.7), and Ca^2+^ event peaks were identified using the same function (‘MinPeakProminence’=0.4).

### SomArchon trace processing to obtain Vm traces in awake mice

We first detrended the raw SomArchon traces by subtracting their two-term exponential fit (MATLAB fit function with ‘exp2’), and then removed the background by subtracting the corresponding rescaled background trace. The rescaled background trace was obtained by first detrending the raw background trace using a two-term exponential model (MATLAB fit function with ‘exp2’), and then rescaling the detrended background trace to match the detrended SomArchon trace using a one-term polynomial model (MATLAB fit function with ‘poly1’). Due to the delay between the SomArchon and GCaMP cameras, we removed the first 50 datapoints from the SomArchon trace to ensure all remaining datapoints were overlapped with the corresponding GCaMP trace. The corresponding SomArchon fluorescence trace was then normalized between the maximum (100%) and minimum (0%) of each trial, so that the percentage represents the dynamic range of the intensity. The resulting trace was considered as membrane voltage (Vm) traces.

### Classification of spikes and detection of slow Vm depolarization in awake mice

Spikes were identified as rapid and large amplitude Vm depolarizations. Specifically, we low-pass filtered the Vm at 10 Hz (f_Vm), and then obtained the bottom half of the f_Vm (f_Vm_low) by replacing f_Vm datapoints that were above its 51-datapoint (∼103 ms) moving average with its moving average. Since the actual Vm baseline fluctuations could be either upward or downward, we estimated the actual standard deviation of Vm baseline fluctuations (fluctuation_std) as double the standard deviation of the f_Vm_low. We then used the MATLAB findpeaks function (‘MinPeakProminence’=5*fluctuation_std) to detect all peaks, and calculated the peak amplitude, defined as the difference between the peak and the lowest datapoint within 5 datapoints (corresponding to ∼10 ms) before and after the spike. The peaks were then clustered using k-means clustering into two groups based on amplitude. We then computed the mean (amp_mean) and standard deviation (amp_std) of the group with larger amplitudes. Spikes were identified as the peaks with amplitude greater than the amp_mean minus two times of the amp_std.

The detected spikes were further classified based on their inter-spike intervals (ISIs) to adjacent spikes and the Vm changes after the spike. Spikes with ISI≥30ms were classified as single spike (SSs), and otherwise bursting spikes (BSs, ISI<30 ms). For each SS, we then identified the occurrence of after-spike-depolarization (ADP), where the averaged Vm during the 5-30 ms time window after the spike was higher than averaged Vm during the 5-30 ms time window before the spike. The chosen time window is consistent with the ADP duration reported previously ^44^. We then classified SSs into those with ADPs (SS-w-ADP) and those without ADPs (SS-wo-ADP). To identify CS, we grouped consecutive BSs with ADPs (here calculated as averaged Vm during 5-60 ms after the spike being higher than the averaged Vm during 5-60 ms before spike), and then identified the onset of CSs as the first BS.

To detect slow Vm depolarization peaks, we performed a 50 data-point (corresponding to ∼100 ms) moving average on Vm traces (mov_Vm), and then used the MATLAB findpeaks function (‘MinPeakProminence’=standard deviation of mov_Vm) across mov_Vm to obtain the depolarization peaks. Due to the limited imaging speed at 500 Hz, in comparison to 10-20 kHz sampling rate typically used for conventional patch clamp recording, spike amplitude appears smaller relative the subthreshold dynamics.

### GCaMP fluorescence trace processing to obtain Ca^2+^ traces, and the identification of Ca^2+^ event peak and rising phase in awake mice

We first removed the background signals by subtracting the raw GCaMP background traces from the raw GCaMP traces, and then detrended by subtracting their two-term exponential fit (MATLAB fit function with ‘exp2’).

Ca^2+^ event peaks and width were then obtained using the MATLAB findpeaks function (‘MinPeakProminence’= standard deviation of the detrended GCaMP trace). The start of a Ca^2+^ event was identified as the local minimum within two event widths prior to the peak using the MATLAB islocalmin function. The rising phase of the Ca^2+^ event was computed as the period between event start and event peak. Ca^2+^ events typically had a rising phase of a couple hundred milliseconds, and thus any rising phase longer than one second was excluded from further analysis. Finally, we upsampled the GCaMP traces to match the corresponding Vm traces, and then normalized by rescaling between the maximum (100%) and the minimum (0%) across all trials, to obtain the Ca^2+^ traces for further analysis.

### Vm and Ca^2+^ traces feature analysis in awake mice

To examine the temporal features of Vm and Ca^2+^ around CS, we first aligned one second of Vm and Ca^2+^ traces to all CS onsets (the first spike within a CS) in a given neuron. We then normalized both traces by subtracting the mean during the 500 ms prior to CS onsets for each neuron, and computed the population mean across all neurons. To avoid the impact of the first spike within a CS on Vm peak identification, we restricted the identification of Vm and Ca^2+^ peak to the 5-500 ms time window after CS onset. Similar analyses were performed to examine Vm and Ca^2+^ temporal features around SS-w-ADP and SS-wo-ADP. To examine the temporal features of Ca^2+^ around slow Vm depolarization peak, Ca^2+^ peaks were identified in the 1-second window centered at Vm depolarization peaks. Similarly, to examine the temporal features of Vm around Ca^2+^ event peaks, Vm peaks were identified in the 1-second window centered at the Ca^2+^ event peaks.

### Characterization of SomArchon and GCaMP8 photobleaching rate

For SomArchon recordings from a neuron, rawF was defined as the mean raw SomArchon fluorescence of the trial, dF was defined as the mean of raw fluorescence of a spike peak minus the minimum raw fluorescence in the 5 ms prior to the spike peak across all spikes detected in each trial, and dF/F was defined as the mean of normalized dF by the raw fluorescence of a spike peak. The time point of each SomArchon trial was estimated by the middle time point of the laser-on period for each snippet trial. The rawF, dF and dF/F of all trials were normalized to those of the first trial from this neuron. Linear regression was conducted between rawF, dF, dF/F, and each trial’s time point respectively. The slope of the linear fitting line indicates the photobleaching rate for each neuron. The photobleaching rates were box plotted for rawF, dF and dF/F, respectively. Two-sample t-test was applied between normal sampling rate group and high sampling group.

For GCaMP recordings from a neuron, rawF was defined as the mean GCaMP8 fluorescence of the trial, dF was defined as the mean of raw fluorescence of a Ca^2+^ event peak minus the raw fluorescence of the starting point of the Ca^2+^ rising phase of the corresponding Ca^2+^ event across all Ca^2+^ events detected in each snippet trial, and dF/F was defined as the mean of normalized dF by the raw fluorescence of a Ca^2+^ peak. The time point of each GCaMP trial was estimated by the middle time point of the blue-LED-on period for each snippet trial. All the other analyses were the same as SomArchon photobleaching characterization.

### Intracranial electrical stimulation in hippocampal CA1 in awake mice

Electrical stimulation was delivered between two intracranial electrodes coupled to the imaging chamber. Electrical pulses were bipolar, square wave, charge balanced, with a pulse width of 100 µs per phase. For 0.7 second sustained electrical stimulations, the pulses were delivered at 40 Hz, 140 Hz, and 1 kHz, controlled by a constant current stimulator (A-M Systems, Model 4100), triggered by TTL pulses generated by MATLAB through a DAQ (National Instruments, BNC-2110 and PCIe-6323). Electrode impedance was approximately 113 kOhm at 40 Hz, 66 kOhm at 140 Hz, 39 kOhm at 1 kHz, measured in the brain between the stimulation electrode and the ground using NanoZ (Plexon, 33377). During each recording, we performed 10 trials while delivering stimulation at a given frequency, with each trial lasting for 3 seconds containing 1 second pre-stimulation baseline, 0.7 second stimulation and 1.3 second post-stimulation. Some neurons were tested with multiple stimulation frequencies. For 1 or 4 pulse electrical stimulation, the inter-pulse interval was set as 15 ms. During each recording, we performed 16 trials while randomizing 1 pulse or 4 pulses for each trial, with each trial lasting for 2 seconds containing 1 second pre-stimulation baseline, 1 pulse (0.2 ms) or 4 pulses (45.2 ms) stimulation and ∼1 second post-stimulation.

To account for variations in surgical preparation across animals, we first empirically determined the stimulation current threshold for each mouse by imaging evoked Vm responses during 40 Hz stimulation. We incrementally increased the current amplitude, starting at 100 µA and increasing by 10-50 µA, until we observed detectable Vm entrainment without noticeable behavioral effects. In subsequent recordings, we started with the current threshold and adjusted the current amplitude by 20-50 µA to ensure clear 40 Hz Vm entrainment without behavioral effects. The same current amplitude was then used for all other neurons recorded and all frequencies applied on the same mouse on the same day. For 0.7 s prolonged stimulation, across all mice, the current amplitude was 100-250 µA (165±62 µA, n=4 recording sessions in 3 mice), corresponding to a charge density of 9.4-23.6 μC/cm^2^ per stimulation phase. For brief one or four-pulse stimulation, the current amplitude was 450-550 µA (500±57.7 µA, n=4 recording sessions in 3 mice), corresponding to a charge density of 42.5-51.9 μC/cm^2^ per stimulation phase.

### Classification of prolonged stimulation evoked Vm and Ca^2+^ responses

To assess stimulation-evoked effects, we first computed the trial-wise mean Vm trace amplitude, mean Ca^2+^ trace amplitude and mean spike rate during stimulation (700 ms) and during the baseline (700 ms immediately prior to each stimulation onset). We assessed significance using the sign-rank test for a pairwise comparison of responses during stimulation versus during baseline (p<0.05). Significantly modulated neurons were further classified as activated or suppressed based on whether the stimulation-evoked change was higher or lower than the baseline respectively.

### Stimulation mediated Vm entrainment evaluation

To determine whether a neuron was significantly entrained, we computed the spike-Vm phase-locking value (PLV) at the stimulation frequency and compared it to a bootstrapped baseline distribution. First, we applied a narrowband filter to the Vm trace at the stimulation frequency (*f*) with a bandwidth of ±10%. We then used a Hilbert transform to find the phase angle (φ) at each timepoint, and calculated the spike-Vm PLV by applying the following equation across all spikes (*N*):

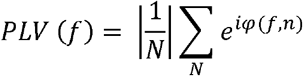

We then built a basal distribution by randomly shuffling the spike onsets and recalculating the PLV 1000 times. Neurons with spike-Vm PLVs higher than the 95^th^ percentile of the bootstrapped distribution were considered significantly entrained at the stimulation frequency.

### Statistical Analysis

All statistical analyses were performed using MATLAB (R2018b, R2021a). One-sample t-test (two-tailed), one-way ANOVA, Wilcoxon rank sum test, and linear fitting were performed using MATLAB built-in functions. For multiple comparisons, we performed Bonferroni correction to determine the p-value threshold for significance. Specifically, for statistical analyses on Ca^2+^ responses associated with SS-wo-ADP and SS-w-ADP, we used 0.0167 (=0.05/3) as the threshold.

## Acknowledgments

We thank members of Han Lab for their help throughout the study, and Jens Christian Schwamborn for technical support on human iPSC derivation. X. H. acknowledges funding from the NIH (1RF1NS129520, 1R01MH122971, 1R01NS115797) and NSF 2002971-DIOS. H.Y.M. acknowledges R01 MH130600. J.M. acknowledges R34NS127098. E.B. acknowledges NSF GRFP DGE-1840990 and NIH T32-EB006359.

## Author contributions

Y.W., H.T. and X.H. designed the study. S.X. and J.M. built the custom wide-field microscope for two-color imaging. Y.W. collected all experimental data. H.T. analyzed the data. Y.W. and E.B. assisted with data analysis. E.B., A.M, H.Y.M and J.C.S assisted with *in vitro* data collection. Y.Z. assisted with *in vivo* data collection. Y.W., H.T. and X.H. prepared the manuscript, and all authors edited the manuscript. X.H. oversaw all aspects of the project and supervised the study.

## Declaration of interests

The authors declare no competing interests.

## Data and materials availability

DNA constructs developed will be deposited to Addgene. Code will be available at the GitHub repository upon manuscript publication. Experimental data are available from the lead contact upon reasonable request.

---

**Supplementary Figure 1.**
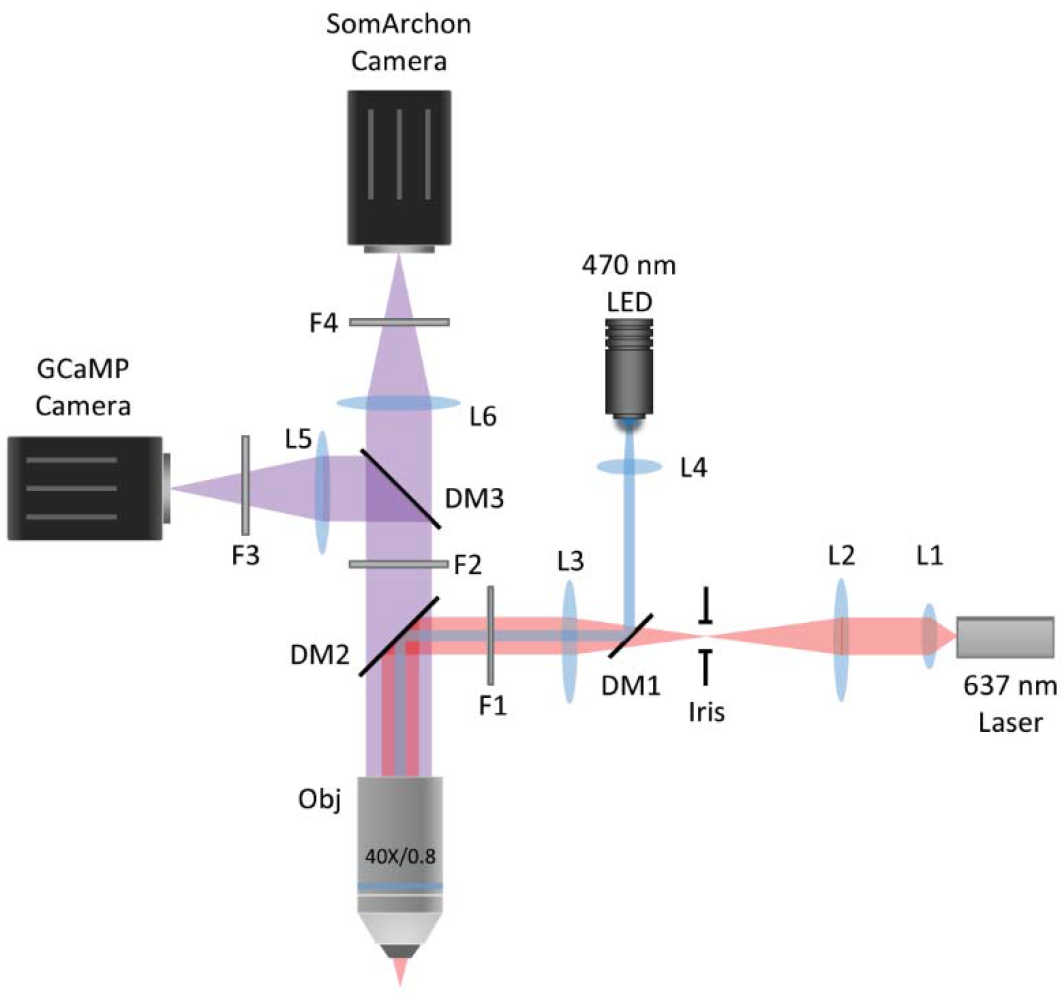
The schematic of the custom widefield microscope for simultaneous imaging of SomArchon and GCaMP fluorescence. L1, high-NA achromatic collimator (Thorlabs F950FC-A). L4, f = 16 mm aspherical condenser (Thorlabs ACL25416U). L2, L3, L5, L6, f = 200 mm achromatic doublet lens (Thorlabs AC254-200-A). DM1, 505 nm long-pass dichromatic mirror (Thorlabs DMLP505R). DM2, quadband dichromatic mirror (Semrock Di03-R405/488/532/635). DM3, multiband dichromatic mirror (Chroma Technology Corp, ZT405/514/635rpc). F1, fluorescence excitation filter (FF01-390/482/532/640-25). F2, fluorescence emission filter (Semrock FF01-446/510/581/703-25). F3, fluorescence emission filter (Semrock FF01-525/45). F4, fluorescence emission filter (Chroma Technology Corp, ET700/75m). Iris, adjustable iris (Thorlabs SM1D12D). 637 nm laser (Ushio America Necsel Red 63x). 470 nm LED (Thorlabs M470L4). SomArchon camera (Hamamatsu Photonics C15440-20UP). GCaMP camera (Hamamatsu Photonics C13440-20CU). Obj, microscope objective (Nikon CFI Apo NIR 40×/0.8NA W).

**Supplementary Figure 2.**
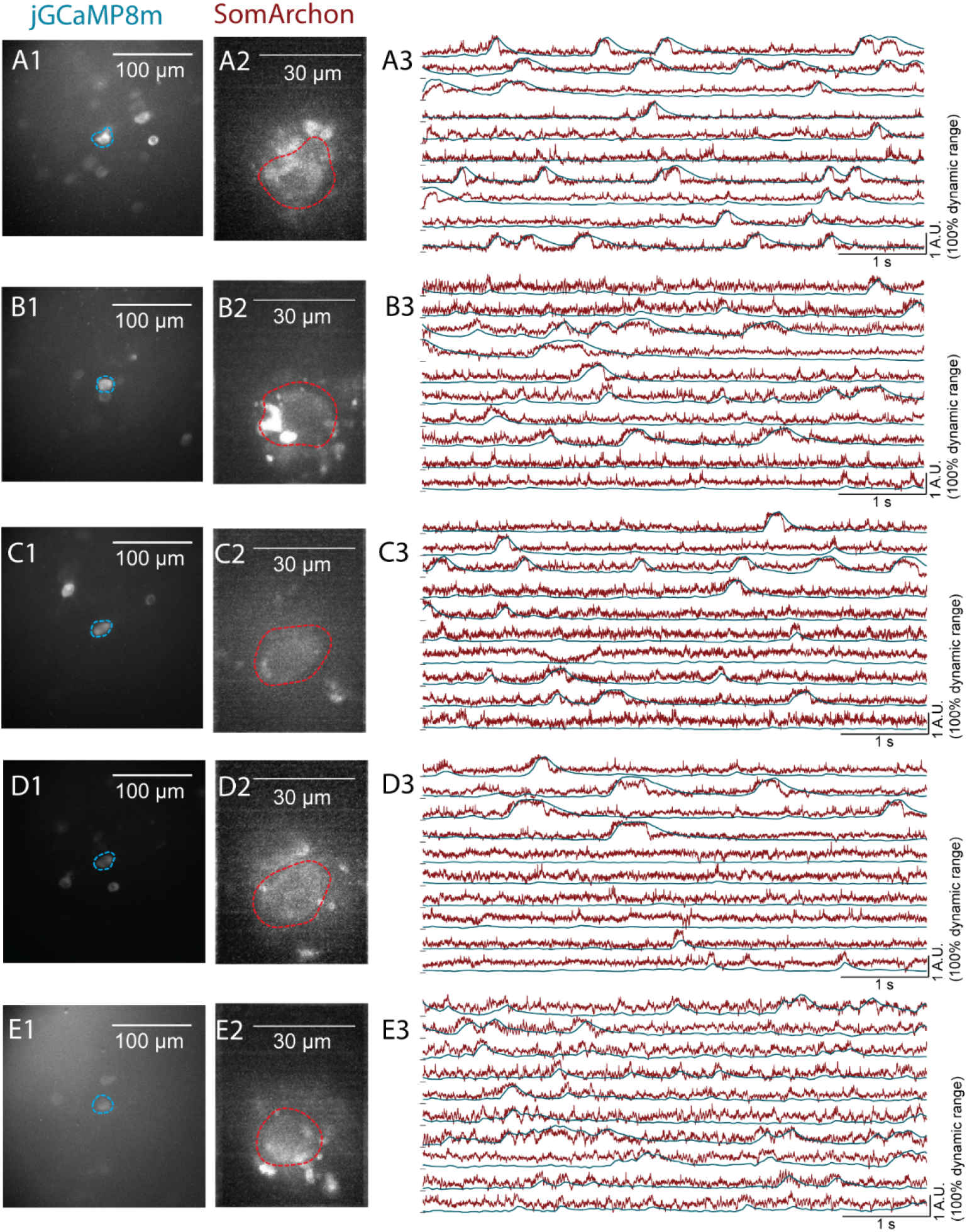
Five example big imaging field of views (fov) in hippocampal CA1 of awake mice, with the centered neuron simultaneously recorded by GCaMP and SomArchon channels. (**A1**) Max-minus-min projection image of a continuous 2-minute GCaMP8 from a big fov from GCaMP channel. The blue outlined neuron had simultaneous SomArchon recordings. (**A2**) Max-minus-min projection image of the centered neuron outlined in (A1) from SomArchon channel. The same neuron was red outlined. (**A3**) The simultaneous Vm (SomArchon) and Ca^2+^ (GCaMP8) traces of the neuron outlined in (A1-A2). (**B-E**) Similar to (A), but for other four CA1 neurons with big GCaMP imaging fovs.

**Supplementary Figure 3.**
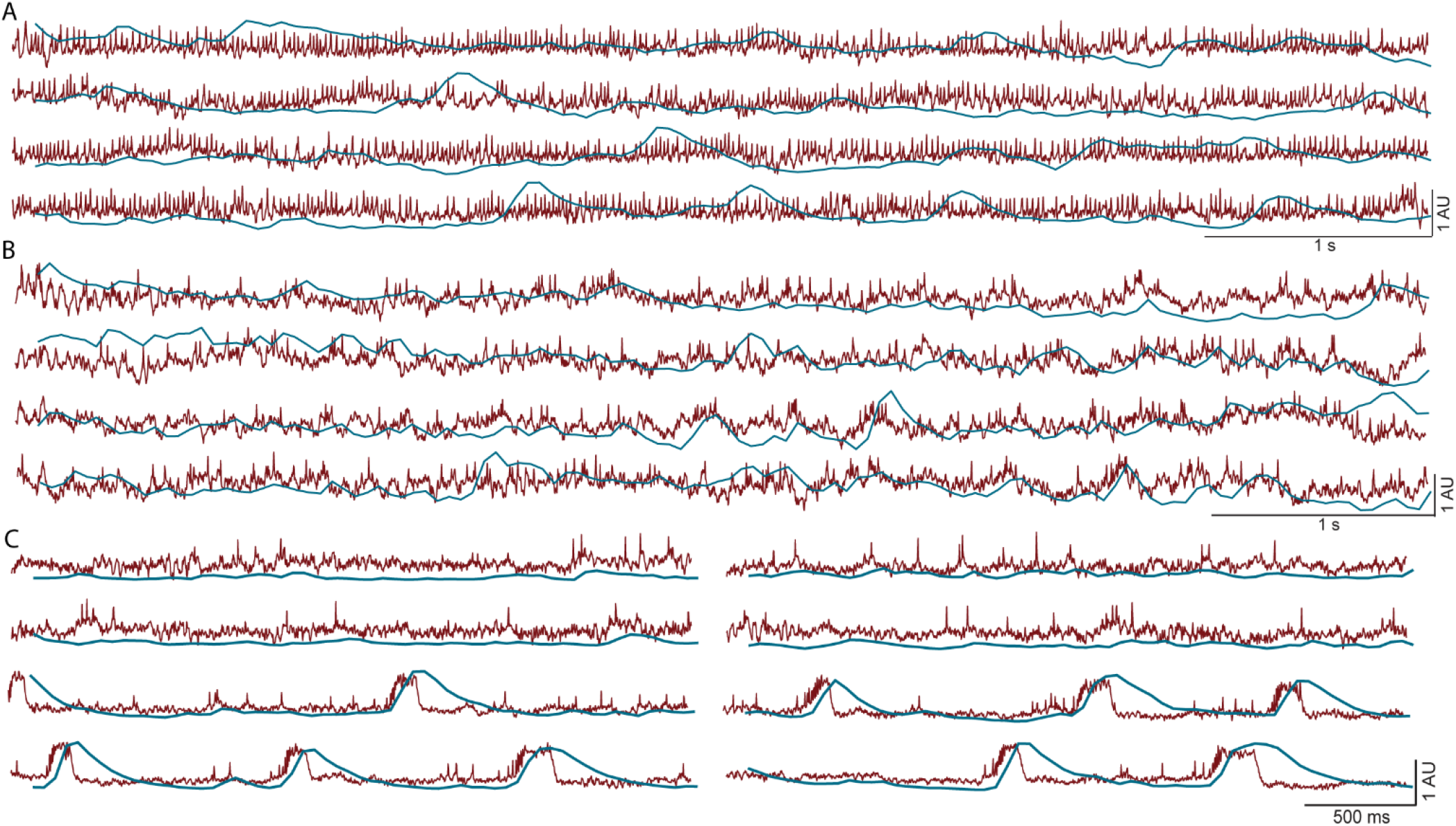
Additional example Vm and Ca^2+^ recordings from CA1 and striatal neurons expressing SomArchon-GCaMP8 in awake mice. (A) A fast spiking CA1 neuron. (B) A fast spiking striatal neuron. (C) A CA1 neuron with transition between predominant SS to CS. Vm is shown in red and Ca^2+^ in blue. 1 AU=100% dynamic range of the continuous recording.

**Supplementary Figure 4.**
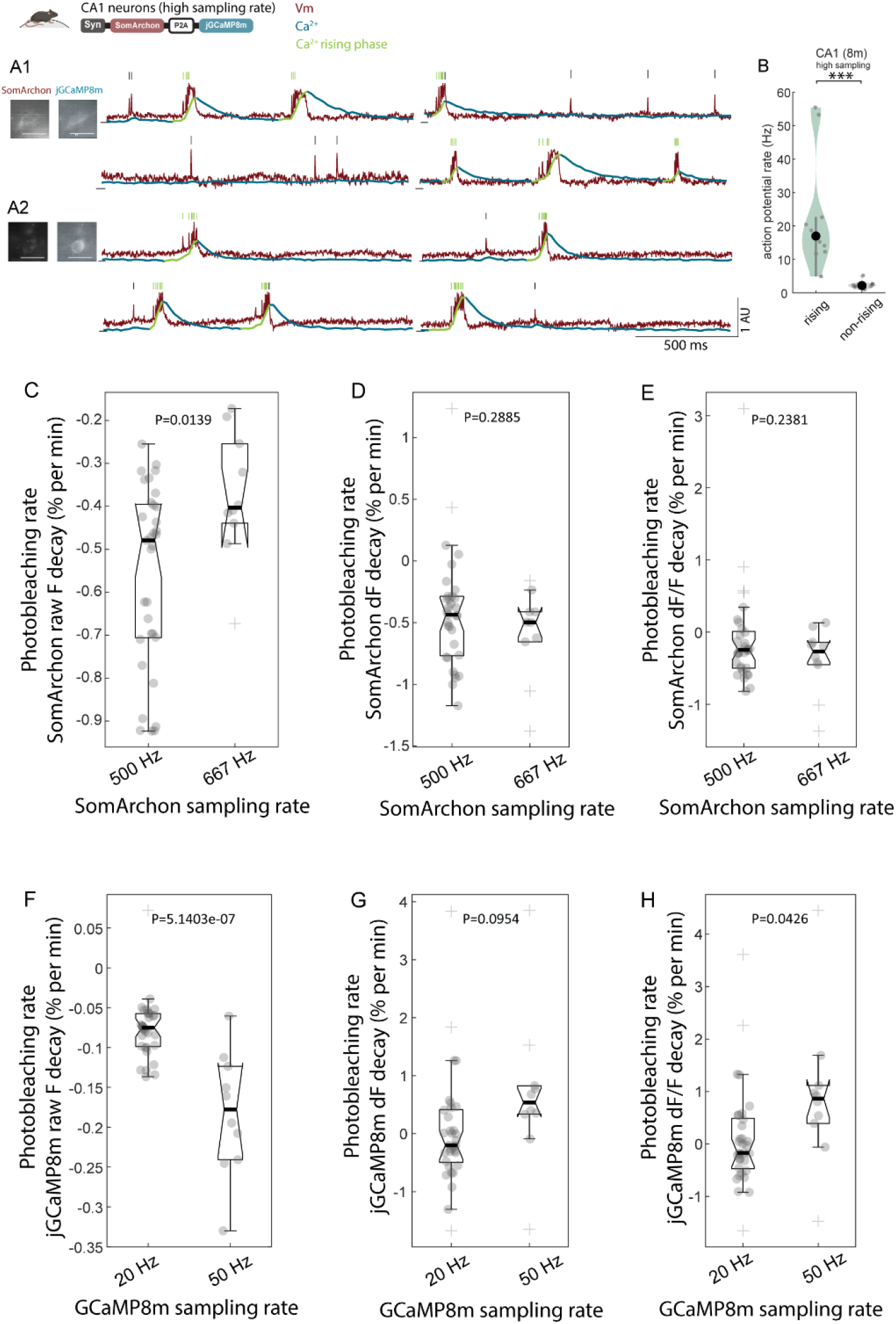
50 Hz Ca^2+^ imaging sampling rate captures more spikes than 20 Hz sampling rate, but photobleaches GCaMP8 faster. (**A1-A2**) Example recordings of Vm (red line) and Ca^2+^ (blue line) from two CA1 neurons in awake mouse brains expressing SomArchon-GCaMP8, showing spikes within (green tick) versus outside (black tick) of Ca^2+^ rising phase (green line overlaid on the blue trace). Scale bar: 30 µm. (**B**) Firing rate during the rising and non-rising phases of Ca^2+^ events. Wilcoxon rank sum test, p=2.5e-4, n=10 neurons. (**C**) Boxplots of the SomArchon raw fluorescence (raw F) decay rate collected at 500 (n=36) versus 667 frames/sec sampling rate (n=10). The central line is the median, notch is 95% confidence interval, and whiskers are the minimum and maximum. Each dot corresponds to a neuron, and + marks outliers. The unpaired t-test p value is displayed in the figure. (**D, E**) Similar to (C), but for SomArchon dF and dF/F decay rate respectively. (**F-H**) Similar to (A-C), but for GCaMP8 photobleaching rates under 20 versus 50 frames/sec. 1 AU=100% dynamic range of the continuous recording.

**Supplementary Figure 5.**
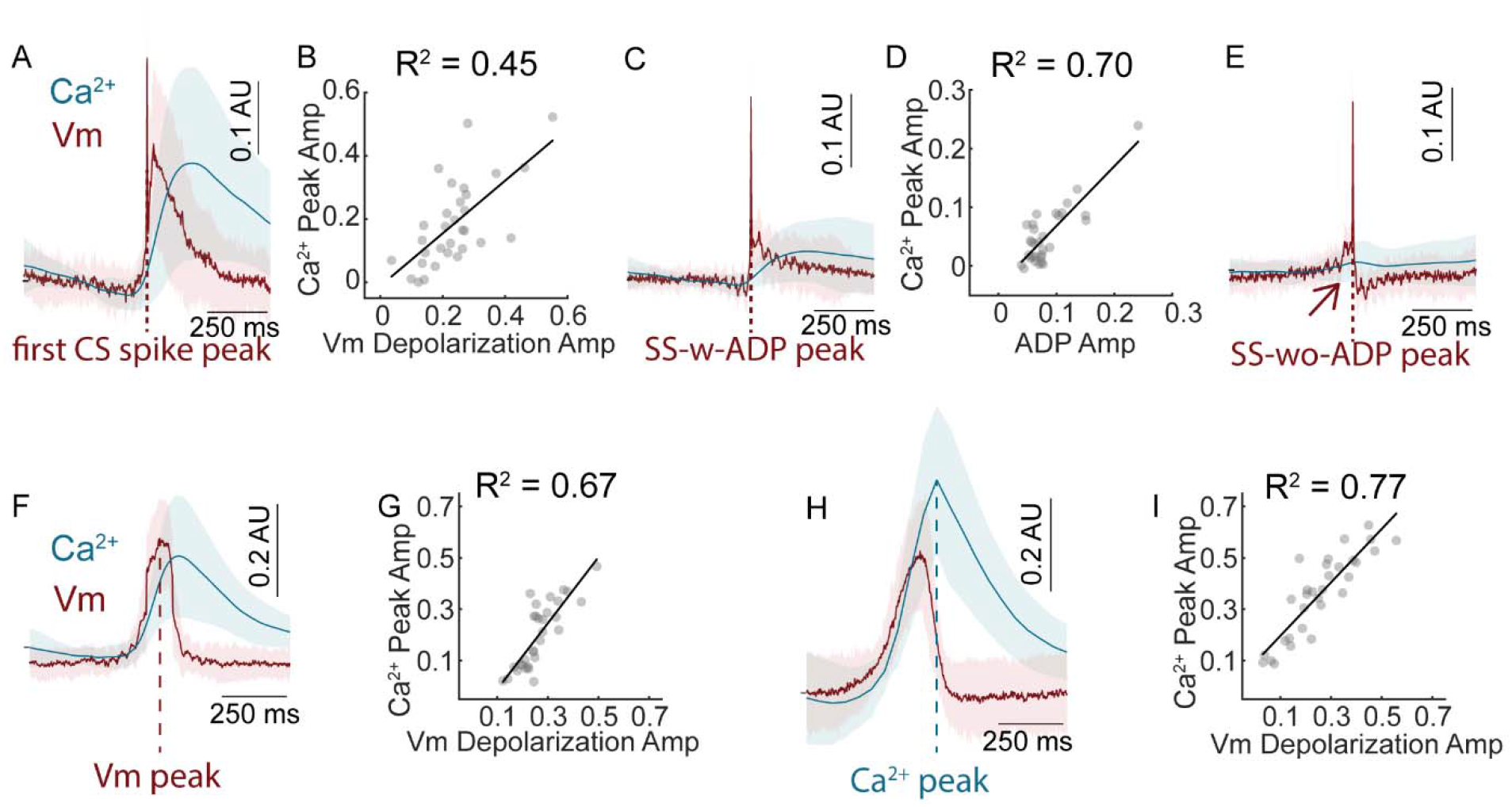
Slow Vm depolarization is tightly correlated with cytosolic Ca^2+^ elevation in CA1 neurons transduced with AA9-SomArchon-P2A-GCaMP7. (**A**) Population Vm (red) and Ca^2+^ (blue) dynamics aligned to CS onset from SomArchon-GCaMP7 expressing neurons (n=31). Shade area: standard deviation. (**B**) Peak amplitude of Vm depolarization versus Ca^2+^ across the 31 neurons. Black line indicates linear regression, with R^2^ value indicated in the figure. (**C, D**) Similar to (A, B), but for SS-w-ADP. (**E**) Similar to (C), but for SS-wo-ADP. (**F**) Similar to (A), but aligned to slow Vm depolarization peaks. (**G**) Peak amplitude of slow Vm depolarization versus Ca^2+^ as shown in (F). (**H, I**) Similar to (F, G), but aligned to Ca^2+^ event peaks. 1 AU=100% dynamic range of the continuous recording.

**Supplementary Figure 6.**
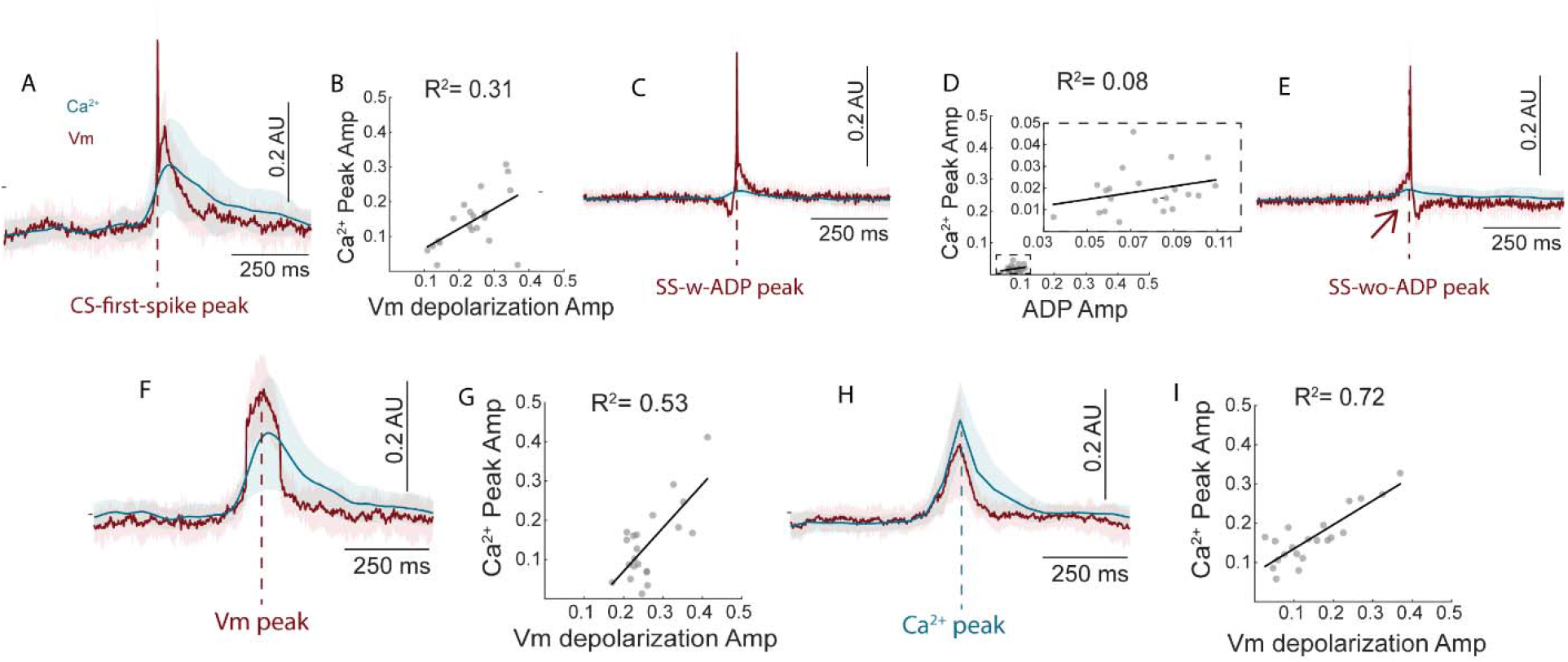
Slow Vm depolarization is tightly correlated with cytosolic Ca^2+^ elevation in dorsal striatal neurons. (**A**) Population Vm (red) and Ca^2+^ (blue) dynamics aligned to first CS spikes from SomArchon-GCaMP8 expressing striatal neurons (n=21). Shade area: standard deviation. (**B**) Peak amplitude of Vm depolarization versus Ca^2+^ across neurons shown in (A) (n=21). Black line indicates linear regression, with R^2^ value indicated in the figure. (**C, D**) Similar to (A, B), but for SS-w-ADP. Since Vm and Ca^2+^ peak amplitudes around SS-w-ADP were small, a zoom-in view is displayed. (**E**) Similar to (C), but for SS-wo-ADP. (**F**) Population Vm and Ca^2+^ dynamics aligned to slow Vm depolarization peaks. (**G**) Peak amplitude of Vm depolarization versus Ca^2+^ (**H, I**) Similar to (F, G), but for Vm and Ca^2+^ dynamics aligned to Ca^2+^ event peaks. 1 AU=100% dynamic range of the continuous recording.

**Supplementary Figure 7.**
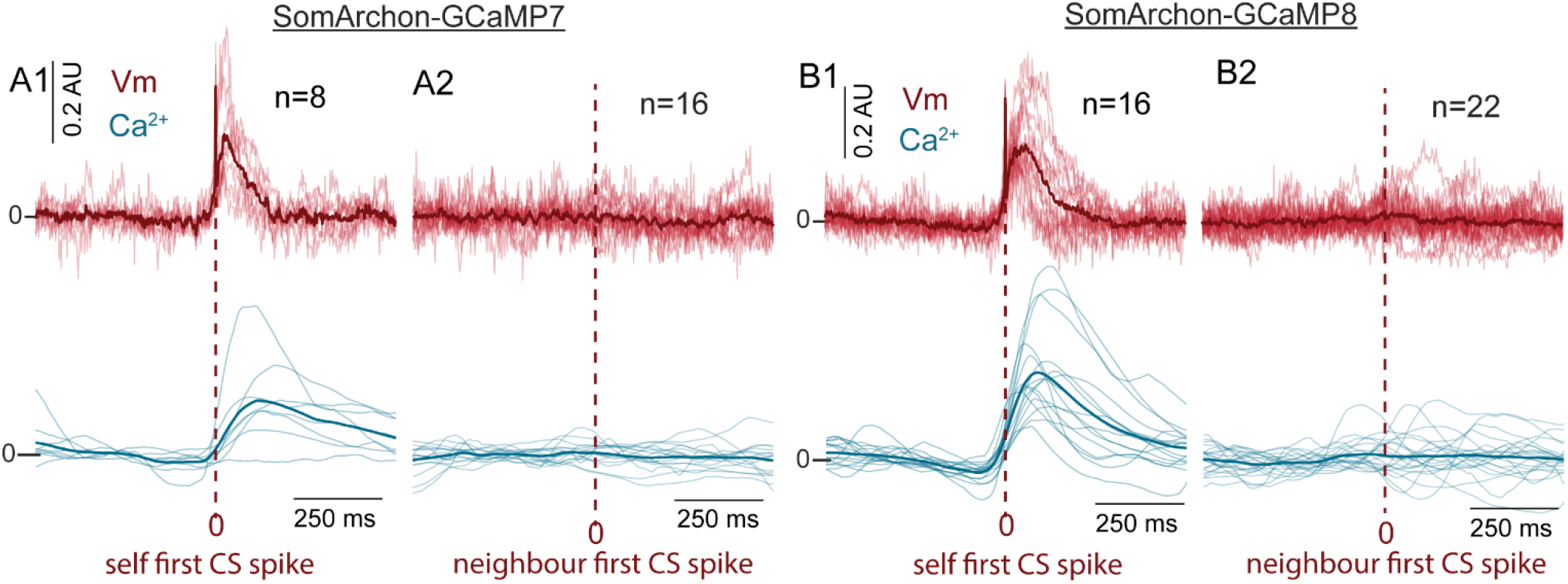
There is little change in Vm and Ca^2+^ when aligned to CS onset of another simultaneously recorded neuron in the same field of view. (**A**) Population Vm (red) and Ca^2+^ (blue) dynamics of one neuron expressing SomArchon-GCaMP7f aligned to CS onset of itself (A1) versus to another simultaneously recorded neuron (A2). Each light pink line indicates one neuron, and the dark red line is the mean. (**B1, B2**) Similar to (A1, A2), but for SomArchon-GCaMP8m expressing neurons. 1 AU=100% dynamic range of the continuous recording.

**Supplementary Figure 8.**
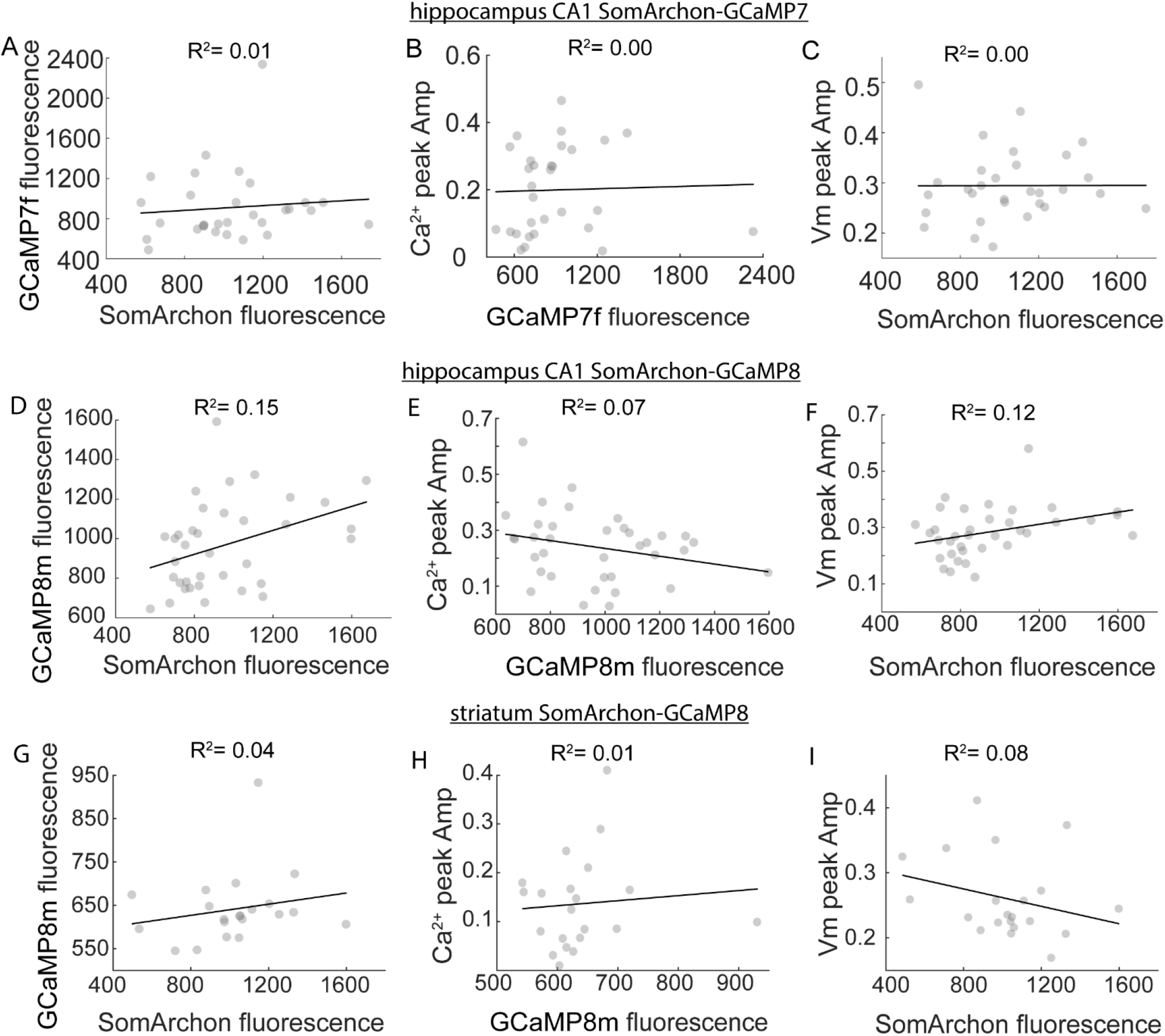
Correlation between basal fluorescence of SomArchon and GCaMP and their peak amplitudes. (**A-C**) CA1 neurons transduced with AAV9-SomArchon-P2A-GCaMP7f (n=31). (A) GCaMP7f basal fluorescence versus SomArchon basal fluorescence in CA1 neurons. (B) Ca^2+^ peak amplitude versus GCaMP7f basal fluorescence. (C) Vm peak amplitude versus SomArchon basal fluorescence. The black lines indicate linear regression, with R^2^ value indicated. **(D-F)** Same as (A-C), but for CA1 neurons transduced by AAV9-SomArchon-P2A-GCaMP8m (n=36). (**G-I**) Same as (A-C), but for striatal neurons transduced with AAV9-SomArchon-P2A-GCaMP8m (n=21).

**Supplementary Figure 9.**
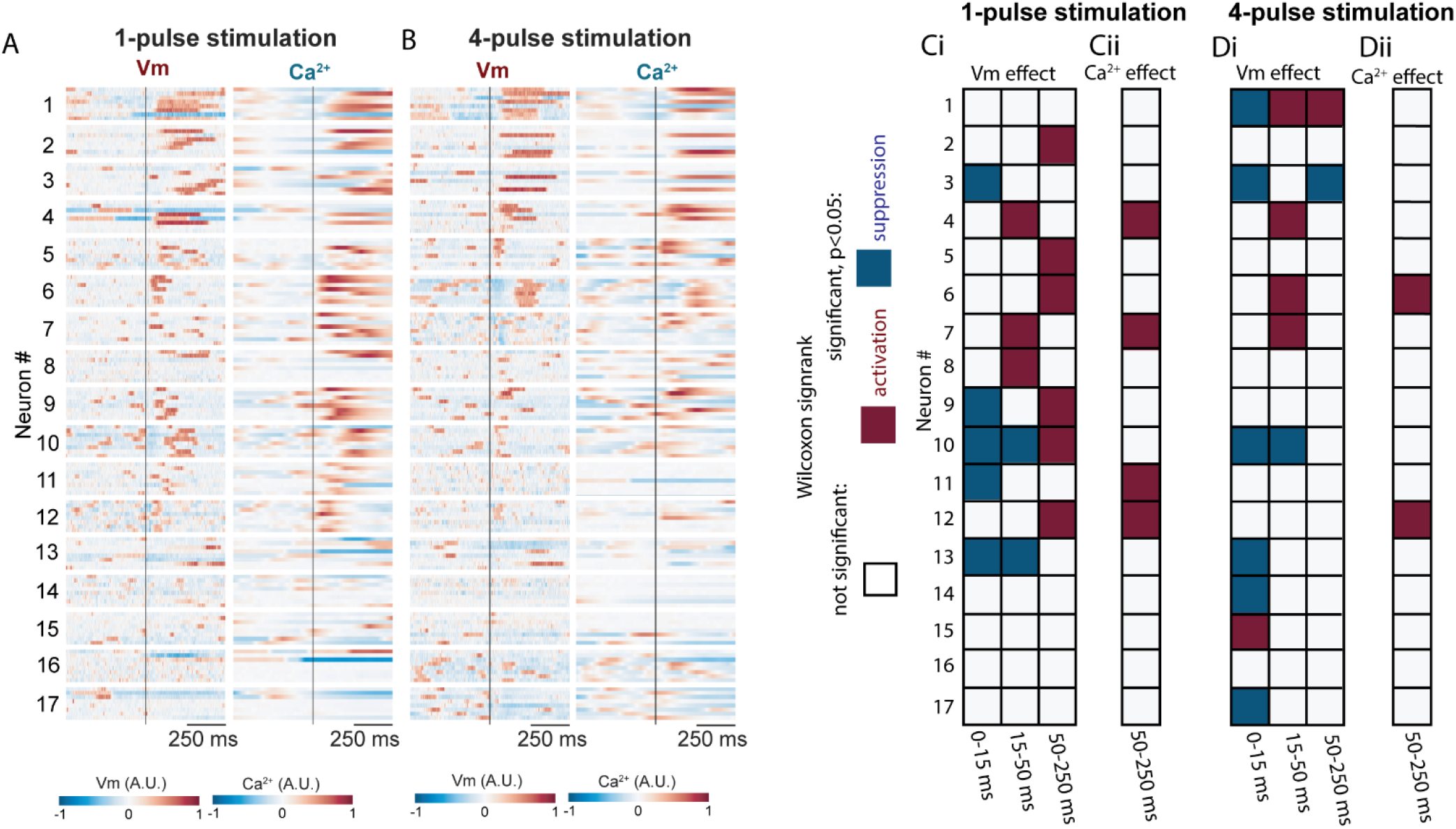
Summary of the brief intracranial electrical stimulation effects on Vm and Ca^2^. (**A**) Heatmap of 1-pulse-stimulation evoked Vm changes across all trials from all neurons, aligned to stimulation onset. Each row corresponds to a trial. Each block corresponds to a neuron and contains 8 trials. Redder color corresponds to stronger Vm depolarization and bluer color corresponds to greater Vm hyperpolarization. Neurons were sorted by mean Vm across 8 trials during the 5-500 ms post the stimulation onset in ascending order. (**B**) Similar to (A), but for the simultaneous Ca^2+^ dynamics from the same neurons with the same sorting order as in (A). (**Ci**) Grid map of 17 neurons’ Vm responses to 1-pulse stimulation across three time windows: 0-15 ms, 15-50 ms, and 50-250 ms after the stimulation onset. Each row corresponds to one neuron, with the same neuron sorting order as in (A). For each neuron and each time window, Wilcoxon signed rank test was conducted across 8 repeated trials. White color indicates no significant difference (p≥0.05) between the examined time window and the corresponding baseline. Blue and red colors indicate significant Vm suppression and activation (p<0.05), respectively. (**Cii**) Similar to (Ci), but for Ca^2+^ responses to the 1-pulse stimulation, with the same neuron sorting order as in (Ci). Due to the limited temporal resolution of 20 Hz GCaMP recording, only 50-250ms time window was examined for Ca^2+^ responses. (**Di-Dii**) Similar to (Ci-Cii), but for 4-pulse stimulation, with the same neuron sorting order as in (Ci).

**Supplementary Figure 10.**
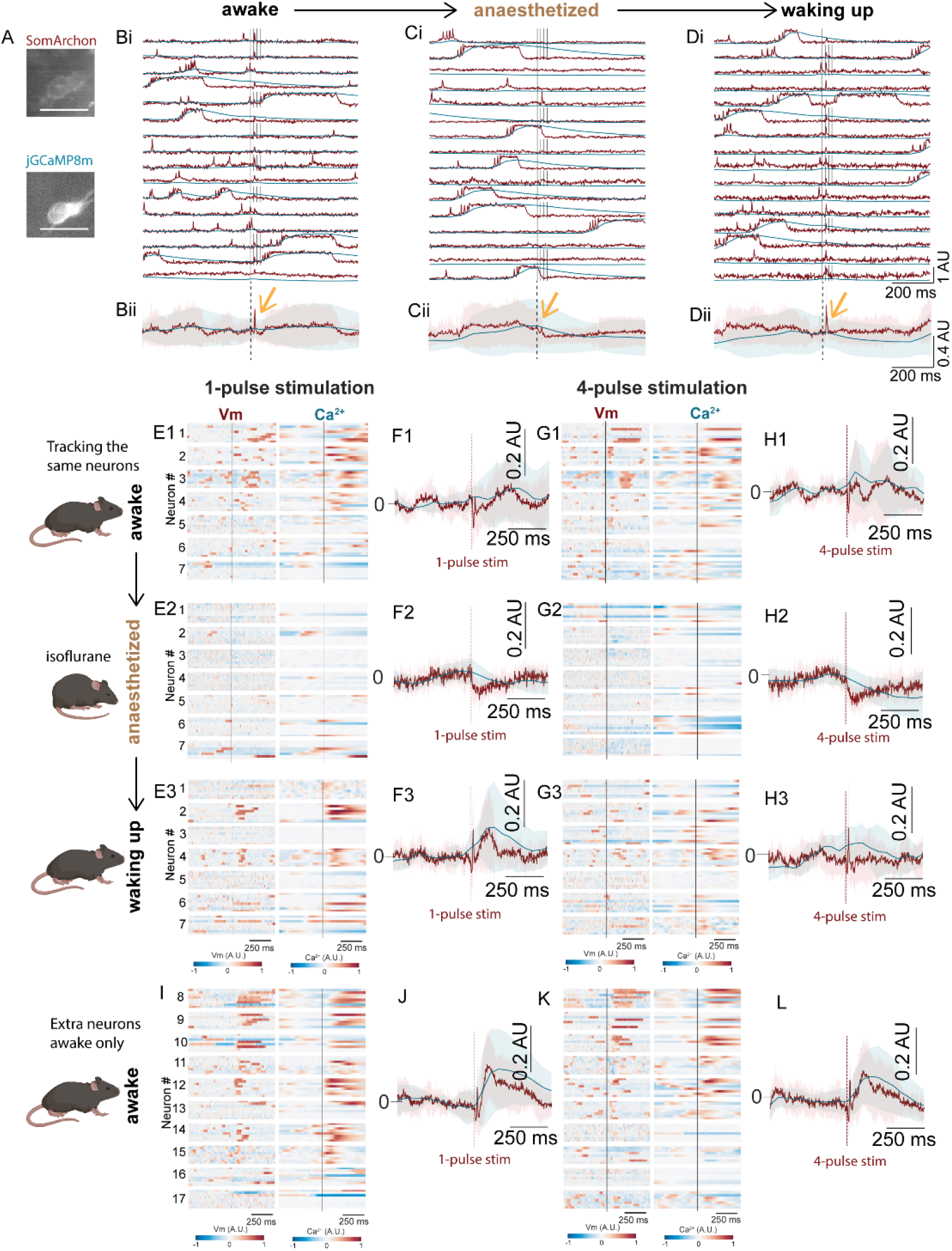
Isoflurane anesthesia reduces neuronal excitability to electrical stimulation. (**A**) The maximum-minus-minimum projection image of SomArchon fluorescence and corresponding GCaMP8m fluorescence of an example CA1 neuron. Scale bars: 30 µm. (**B-D**) Vm and Ca^2+^ traces recorded in the neuron shown in (A) when the mouse was awake (Bi), under anesthesia (Ci), and post-anesthesia awake at 40 minutes after isoflurane offset (Di). The onset of each electrical stimulation pulse is indicated by the black lines. (Bii-Dii) Mean evoked response across trials shown in (Ci-Ei). Shaded areas correspond to standard deviation. (**E1-E3**) Heatmaps of 1-pulse-evoked Vm (left) and Ca^2+^ traces (right) from 7 neurons recorded across awake, under-anesthesia, and post-aneasthsia awake states. Each row corresponds to one trial, and each block corresponds to 8 trials for a neuron. Redder color corresponds to stronger Vm depolarization and Ca^2+^ increase; bluer color corresponds to greater Vm hyperpolarization and Ca^2+^ decrease. (**G1-G3**) Similar to (E1-E3), but for 4-pulse-evoked Vm and Ca^2+^ traces from the same 7 neurons, and with the same neuron order. (**F1-F3**) Population Vm (red) and Ca^2+^ (blue) dynamics aligned to the onset of 1-pulse stimulation from the neurons in (E1-E3). Shade area: standard deviation. (**H1-H3**) Similar to (F1-F3), but for 4-pulse stimulation. (**I**, **J**, **K**, **L**) Similar to (E1, F1, G1, H1), but from 10 other CA1 neurons recorded only during the awake state. 1 AU=100% dynamic range of the continuous recording.

**Supplementary Figure 11.**
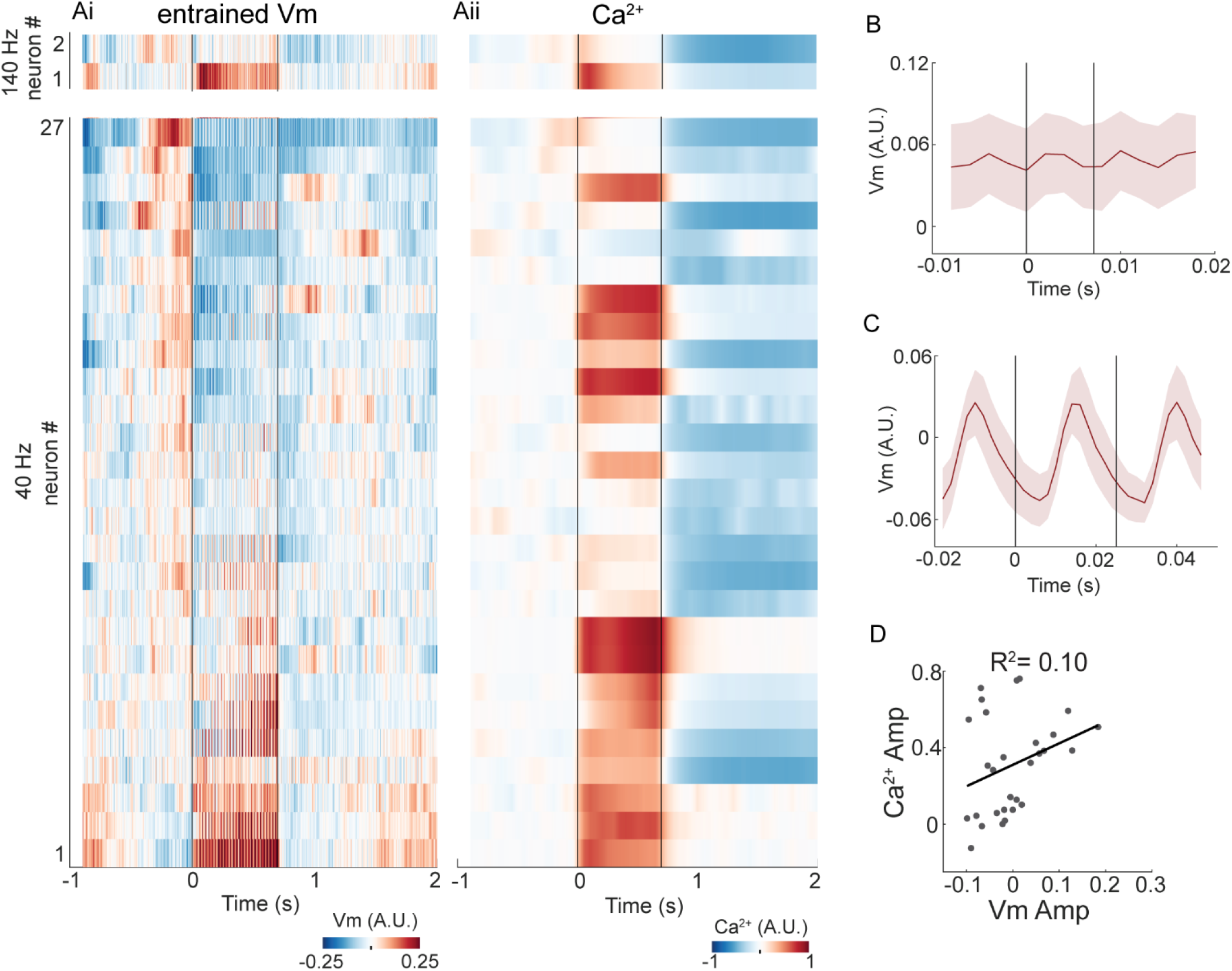
Vm and Ca^2+^ dynamics in Vm-entrained neurons during prolonged intracranial stimulation. (**Ai**) Heatmaps of stimulation-evoked Vm changes across neurons entrained by 40Hz stimulation (n=27 neurons, out of 29 recorded), and 140Hz stimulation (n=2 neurons, out of 20 recorded). Redder color corresponds to stronger Vm depolarization and bluer color corresponds to greater Vm hyperpolarization. Within each frequency group, neurons were sorted by mean Vm changes during the stimulation period in ascending order. (**Aii**) Corresponding heatmaps of evoked Ca^2+^ changes, with the same neuron sorting order as the Vm heatmap in (Ai). (**B**) Vm changes aligned to individual stimulation pulses during 140Hz stimulation (n=2 neurons, and 98 pulses per neuron). Solid line is the mean across all pulses and all neurons, and shaded area corresponds to standard deviations. (**C**) Same as (B), but during 40Hz stimulation (n=27 neurons, and 28 pulses per neuron). (**D**) The mean amplitude of stimulation-evoked Vm versus Ca^2+^ across all Vm-entrained neurons (n=29). Black line indicates linear regression (R^2^=0.10, p=0.104). 1 A.U.=100% dynamic range of the continuous recording.

**Supplementary Figure 12.**
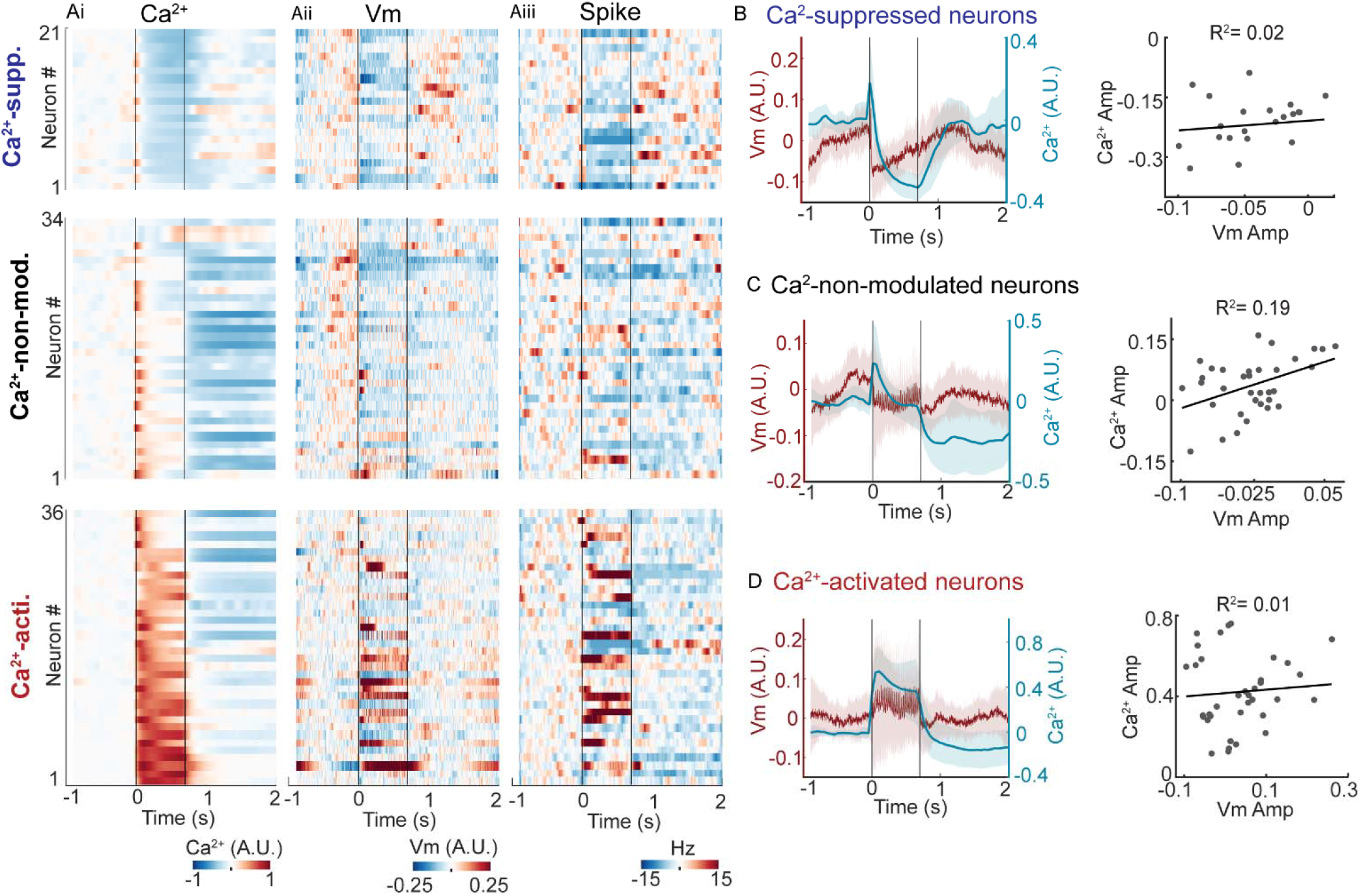
Prolonged intracranial electrical stimulation effect on Vm and Ca^2+^ across neuron populations categorized based on evoked Ca^2+^ changes. (**Ai**) Heatmaps of stimulation-evoked Ca^2+^ changes, aligned to stimulation onset, across neurons that were suppressed (top), non-modulated (middle) and activated (bottom). Each row corresponds to a neuron. Redder color corresponds to stronger Ca^2+^ increase and bluer color corresponds to greater Ca^2+^ decrease. Within each group, neurons were sorted by mean Ca^2+^ changes during the stimulation period in ascending order. Note, the heatmap is a resorted version of the heatmap shown in Fig. 5Aii and Fig. 5Eiii. (**Aii, Aiii**) Corresponding heatmaps of evoked Vm changes (Aii) and spike rate changes (Aiii), with the same neuron sorting order as in (Ai). (**B**) Left: Mean evoked Vm (red) and Ca^2+^ (blue) across Ca^2+^-suppressed neurons. Shade areas indicate standard deviations. Right: The mean amplitude of evoked Vm versus Ca^2+^ in Ca^2+^-suppressed neurons (n=21 neurons, R^2^=0.02, p=0.578). Line indicates linear regression with the R^2^ indicated on the top. (**C**) Similar to (B), but for Ca^2+^-non-modulated neurons (n=34 neurons, R^2^=0.19, p=0.01). (**D**) Similar to (B), but for Ca^2+^-activated neurons (n=36 neurons, R^2^=0.01, p=0.648). 1 A.U.=100% dynamic range of the continuous recording.

**Table S1.**
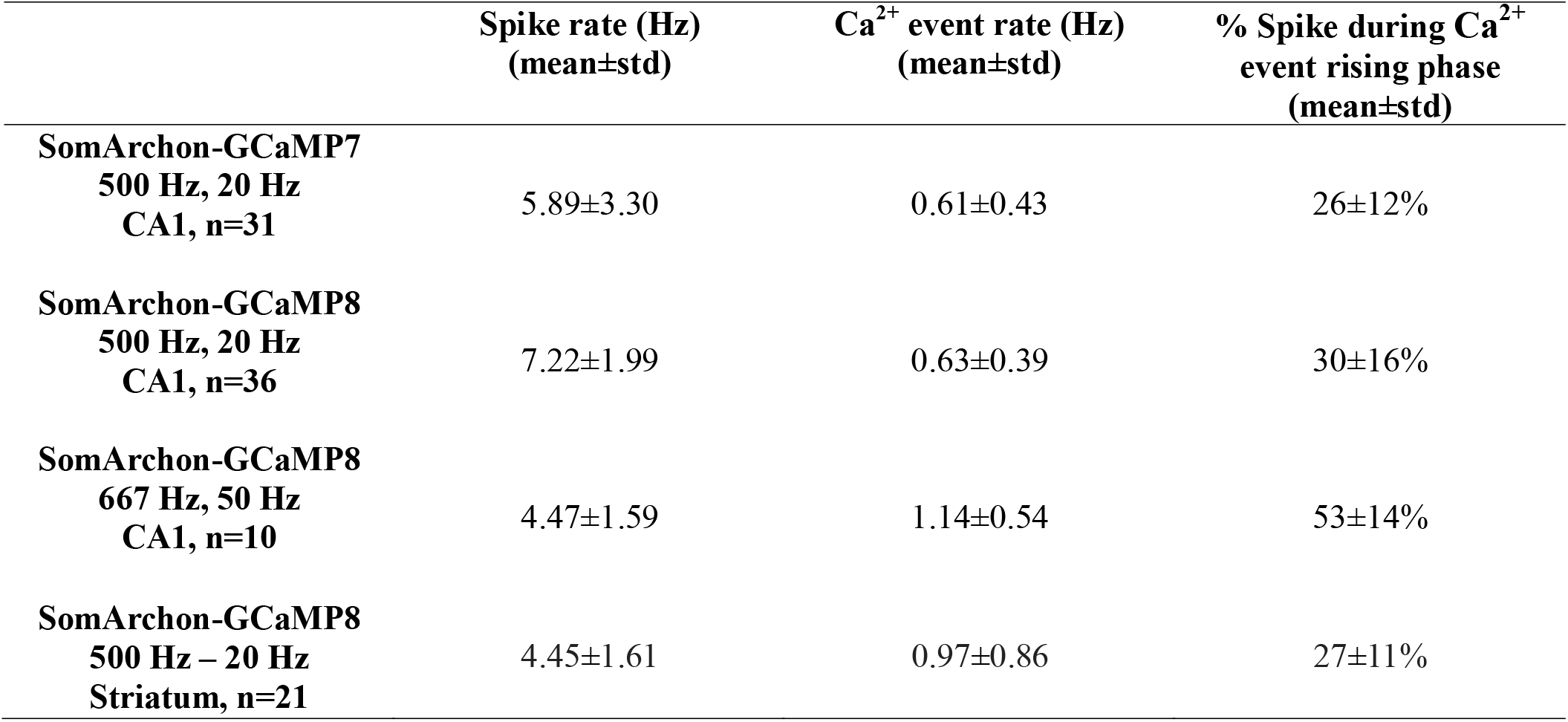
Summary of spike rate, Ca^2+^ event rate, and % of spikes during the rising phase of Ca^2+^ events. Left column: Name of the sensor, imaging sampling rate of SomArchon and GCaMP, recorded brain region, and n indicating total neuron numbers in each group.

**Table S2.** Summary of prolonged electrical stimulation effects on Vm, spike rate, and Ca^2+^. Supp., suppressed. Non-mod., non-modulated. Acti., Activated. N=91 neurons over all electrical stimulation frequencies, including N=29 neurons for 40Hz, N=20 neurons for 140Hz and N=42 neurons for 1kHz.

| Neural activity | Stimulation Effect | Mixed frequencies |  | Specific frequency |  | Ca<sup>2+</sup> |  |  |
| --- | --- | --- | --- | --- | --- | --- | --- | --- |
|  |  | Modulated/total | Stimulation frequency | Modulated/total |  | Supp. | Non-mod. | Acti. |
| Vm | Supp. | 40/91 | 40Hz | 13/29 |  | 0/29 | 6/29 | 7/29 |
|  |  |  | 140Hz | 9/20 |  | 0/20 | 7/20 | 2/20 |
|  |  |  | 1kHz | 18/42 |  | 12/42 | 3/42 | 3/42 |
|  | Non-mod. | 32/91 | 40Hz | 9/29 |  | 2/29 | 4/29 | 3/29 |
|  |  |  | 140Hz | 4/20 |  | 0/20 | 2/20 | 2/20 |
|  |  |  | 1kHz | 19/42 |  | 7/42 | 10/42 | 2/42 |
|  | Acti. | 19/91 | 40Hz | 7/29 |  | 0/29 | 0/29 | 7/29 |
|  |  |  | 140Hz | 7/20 |  | 0/20 | 2/20 | 5/20 |
|  |  |  | 1kHz | 5/42 |  | 0/42 | 0/42 | 5/42 |
| Spike | Supp. | 21/91 | 40Hz | 7/29 |  |  |  |  |
|  |  |  | 140Hz | 6/20 |  |  |  |  |
|  |  |  | 1kHz | 8/42 |  |  |  |  |
|  | Non-mod. | 56/91 | 40Hz | 15/29 |  |  |  |  |
|  |  |  | 140Hz | 10/20 |  |  |  |  |
|  |  |  | 1kHz | 31/42 |  |  |  |  |
|  | Acti. | 14/91 | 40Hz | 7/29 |  |  |  |  |
|  |  |  | 140Hz | 4/20 |  |  |  |  |
|  |  |  | 1kHz | 3/42 |  |  |  |  |
| Ca<sup>2+</sup> | Supp. | 21/91 | 40Hz | 2/29 |  |  |  |  |
|  |  |  | 140Hz | 0/20 |  |  |  |  |
|  |  |  | 1kHz | 19/42 |  |  |  |  |
|  | Non-mod. | 34/91 | 40Hz | 10/29 |  |  |  |  |
|  |  |  | 140Hz | 11/20 |  |  |  |  |
|  |  |  | 1kHz | 13/42 |  |  |  |  |
|  | Acti. | 36/91 | 40Hz | 17/29 |  |  |  |  |
|  |  |  | 140Hz | 9/20 |  |  |  |  |
|  |  |  | 1kHz | 10/42 |  |  |  |  |

## Notes

### Competing Interest Statement

The authors have declared no competing interest.

### Summary of Updates

We revised the technical details and related results, and updated the author list.

